# Comparative Analysis of Human-Chimpanzee Divergence in Brain Connectivity and its Genetic Correlates

**DOI:** 10.1101/2024.06.03.597252

**Authors:** Yufan Wang, Luqi Cheng, Deying Li, Yuheng Lu, Changshuo Wang, Yaping Wang, Chaohong Gao, Haiyan Wang, Wim Vanduffel, William D. Hopkins, Chet C. Sherwood, Tianzi Jiang, Congying Chu, Lingzhong Fan

**Author notes:** These authors contributed equally (Yufan Wang, Luqi Cheng). **Corresponding Authors:** Lingzhong Fan, Institute of Automation, Chinese Academy of Sciences, Beijing 100190, China., or Congying Chu, Institute of Automation, Chinese Academy of Sciences, Beijing 100190, China.

## Abstract

Chimpanzees (*Pan troglodytes*) are humans’ closest living relatives, making them the most directly relevant comparison point for understanding human brain evolution. Zeroing in on the differences in brain connectivity between humans and chimpanzees can provide key insights into the specific evolutionary changes that might have occured along the human lineage. However, conducting comparisons of brain connectivity between humans and chimpanzees remains challenging, as cross-species brain atlases established within the same framework are currently lacking. Without the availability of cross-species brain atlases, the region-wise connectivity patterns between humans and chimpanzees cannot be directly compared. To address this gap, we built the first Chimpanzee Brainnetome Atlas (ChimpBNA) by following a well-established connectivity-based parcellation framework. Leveraging this new resource, we found substantial divergence in connectivity patterns across most association cortices, notably in the lateral temporal and dorsolateral prefrontal cortex between the two species. Intriguingly, these patterns significantly deviate from the patterns of cortical expansion observed in humans compared to chimpanzees. Additionally, we identified regions displaying connectional asymmetries that differed between species, likely resulting from evolutionary divergence. Genes associated with these divergent connectivities were found to be enriched in cell types crucial for cortical projection circuits and synapse formation. These genes exhibited more pronounced differences in expression patterns in regions with higher connectivity divergence, suggesting a potential foundation for brain connectivity evolution. Therefore, our study not only provides a fine-scale brain atlas of chimpanzees but also highlights the connectivity divergence between humans and chimpanzees in a more rigorous and comparative manner and suggests potential genetic correlates for the observed divergence in brain connectivity patterns between the two species. This can help us better understand the origins and development of uniquely human cognitive capabilities.

## Introduction

Chimpanzees (*Pan troglodytes*) are among humans’ (*Homo sapiens*) closest living primate relatives, with a shared ancestor dating back approximately 6-8 million years ago ^1^. Although chimpanzees have brains that are approximately one-third the brain size of humans ^2,3^, they demonstrate similarities in neuroanatomical structure ^4,5^ and cognitive functions ^6–9^, including social behavior, working memory, and tool use. Given their greater genetic proximity and more comparable neurobiology to humans than other primates ^10^, elucidating chimpanzee brain organization is critical as a comparative reference for understanding human evolution. Neuroimaging has enabled quantitative comparisons of brain structure between chimpanzees and other primates ^5,11–13^. However, morphological changes in cortical areas alone cannot fully explain evolutionary adaptions, especially with regard to the association cortices ^14,15^.

Evolutionary changes in wiring space, especially in the white matter tracts beneath the cortex, significantly influence anatomical and functional differences between species ^14,16,17^. These anatomical connections characterize brain regions and support flexible cognitive functions ^18,19^. Recent studies have mapped chimpanzee brain connections using diffusion MRI and revealed substantial interspecies differences with humans ^14,16,17,20–24^. Moreover, understanding brain evolution requires a genetic perspective, as genetic factors strongly influence neural connectivity and uncover molecular mechanisms driving interspecies differences in brain connections, revealing insights into cognitive diversity and adaptation among primates ^25,26^. However, comprehensive whole-brain connectional analyses and comparisons between chimpanzees and humans, coupled with genetic investigations into species differences in brain connections, are still lacking. Another critical consideration is the neuroanatomical and functional asymmetries observed in both human and non-human primates ^27,28^. These asymmetries have been linked to faculties that require higher-order cognitive processing, such as language and complex tool use ^29,30^. Previous comparative asymmetry studies mainly examined local structural features ^31,32^, and limited research on hemispheric asymmetries of connections to date only focused on few localized regions ^33,34^. Therefore, mapping brain-wide connectional asymmetries could illuminate lateralized human-specific cognitive functions.

However, a major challenge in cross-species neuroscience is the lack of a common brain reference system that facilitates comparison across different species. Therefore, the development of brain atlases capable of facilitating such comparisons is essential for revealing the (dis)similarities between species at the level of biologically meaningful subregions ^35^. Previous comparative analyses defined homologous brain regions between species using cytoarchitecture, myeloarchitecture, macroanatomy, connectivity patterns, functional activation, or a combination of these features, leading to a variety of brain atlases for humans ^36–39^ and chimpanzees ^12,21,40,41^. However, inconsistencies in the modalities and scales used to construct these atlases render cross-species comparisons challenging. Recent connectivity-based parcellation successfully delineated distinct brain areas in humans, macaques, and marmosets using anatomical connections ^37,42,43^. This demonstrates the feasibility of parcellating the chimpanzee brain based on diffusion MRI. Such a standardized atlasing approach also enables meaningful cross-species comparisons ^33^. Meanwhile, generating homologous white matter tracts across species within a connectivity blueprint framework has been used to predict homologous areas between species, even when their relative locations have changed. This approach also helps identify unique aspects of brain organization, offering new opportunities for investigating evolutionary changes in brain wiring ^44,45^.

To comprehensively investigate chimpanzee-human connectional divergence, in the present study, we first created the most refined atlas of the chimpanzee brain, i.e., the Chimpanzee Brainnetome Atlas (ChimpBNA), following a well-established connectivity-based parcellation framework ^37^ (Figure 1A). We then reconstructed homologous white matter tracts and built connectivity blueprints for both humans and chimpanzees ^44,45^. Leveraging these blueprints, we investigated cross-species connectivity divergence at the subregion level and their associated white matter tracts (Figure 1B). Next, we examined the lateralization of connectivity patterns across the entire brain in each species and aligned it to a common space ^14^ to explore the relationship to connectivity divergence between species (Figure 1C). Finally, we identified the genes and their expression patterns associated with connectivity divergence between the species (Figure 1D). This fine-grained chimpanzee parcellation and comprehensive cross-species connectional analysis provide new insights into human brain evolution.

**Figure 1.**
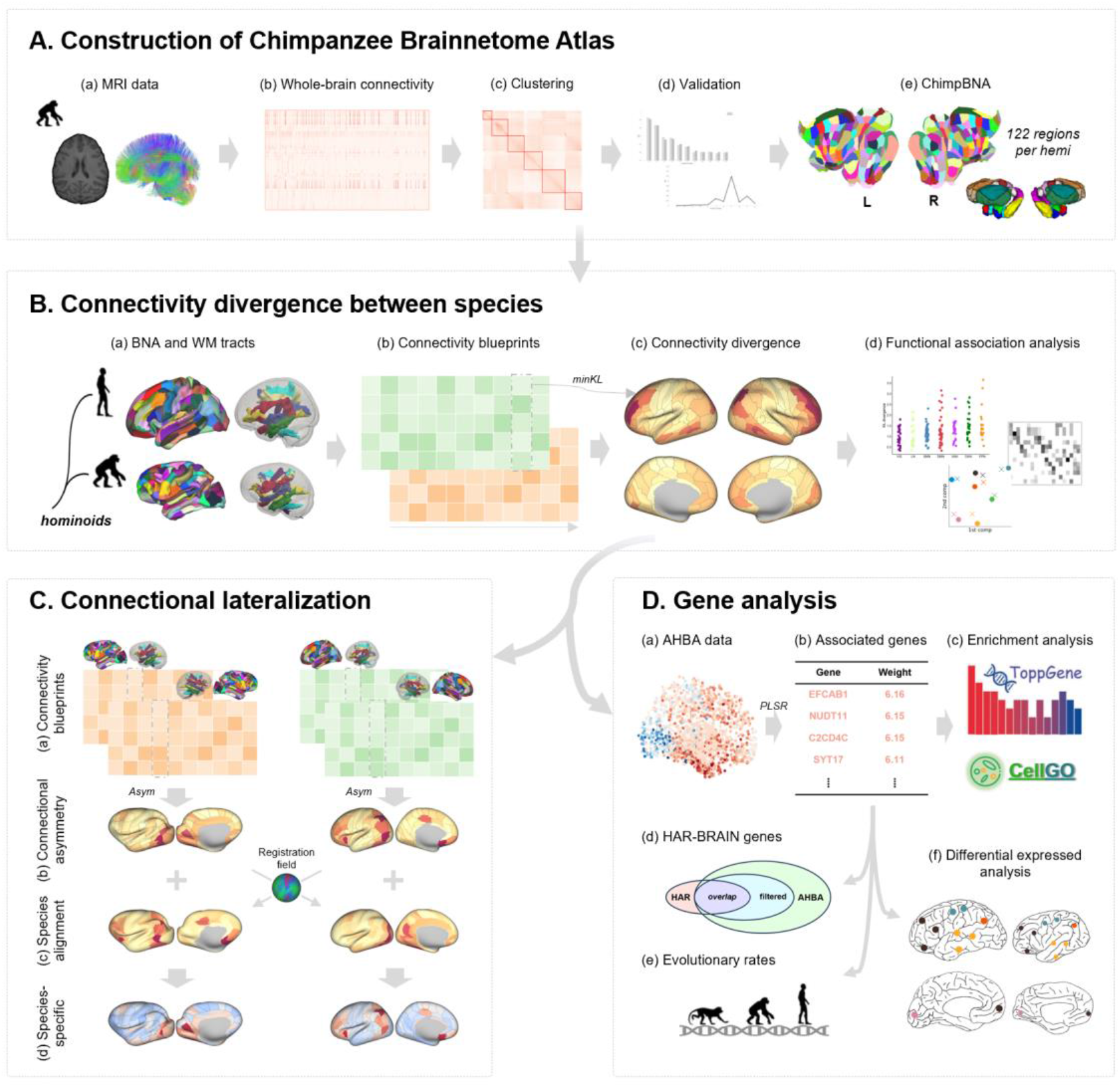
Analysis pipeline. **(A)** Following a connectivity-based parcellation procedure, we used MRI data from chimpanzee brains (a) to construct the Chimpanzee Brainnetome Atlas. Tractography and similarity matrices were performed (b) for subsequent spectral clustering (c). The clustering results were validated using several indices (d), and the final parcellation of the whole brain was obtained (e). **(B)** We utilized the Brainnetome Atlas of humans and chimpanzees and homologous white matter tracts (a), to build regional connectivity blueprints for each species (b). The blueprints were used to explore the connectivity divergence between humans and chimpanzees (c), followed by functional association analysis of this divergence (d). **(C)** We used the connectivity blueprints from two hemispheres for each species (a) to investigate the asymmetric connectivity pattern (b). Myelin-based registration was used to align the two species into a common space (c). Thus the species-specific asymmetric connectivity pattern could be calculated (d). **(D)** AHBA data (a) was used to identify the genes associated with the connectivity divergence by PLSR (b), the filtered genes were input to gene enrichment analysis and cell-type enrichment analysis (c), as well as evolutionary investigation, including overlap with HAR-BRAIN genes (d), evolutionary rates (e), and differentially expressed analysis between the two species (f).

## Results

### Connectivity-based Parcellation of the Chimpanzee Brain

Following the connectivity-based parcellation framework modified from our previous study (Figure S1) ^37^, we first delineated 26 initial seed masks (19 cortical and 7 subcortical masks) on the chimpanzee brain template and registered them to individual brains. We calculated the whole-brain connectivity of each region and subdivided it into several clusters following the validation of indices based on the reproducibility of the subjects’ interhemispheric consistency in the topological relationships. Using the connectivity derived from the probabilistic tractography of 46 chimpanzees *in vivo* diffusion MRI data, we subdivided the brain into 200 cortical (Figure 2A) and 44 subcortical regions (Figure S2), thereby building ChimpBNA, the most refined atlas of the chimpanzee brain to date. The ChimpBNA parcellation is interactively accessible via the web viewer (Figure S3, https://molicaca.github.io/atlas/chimp_atlas.html).

**Figure 2.**
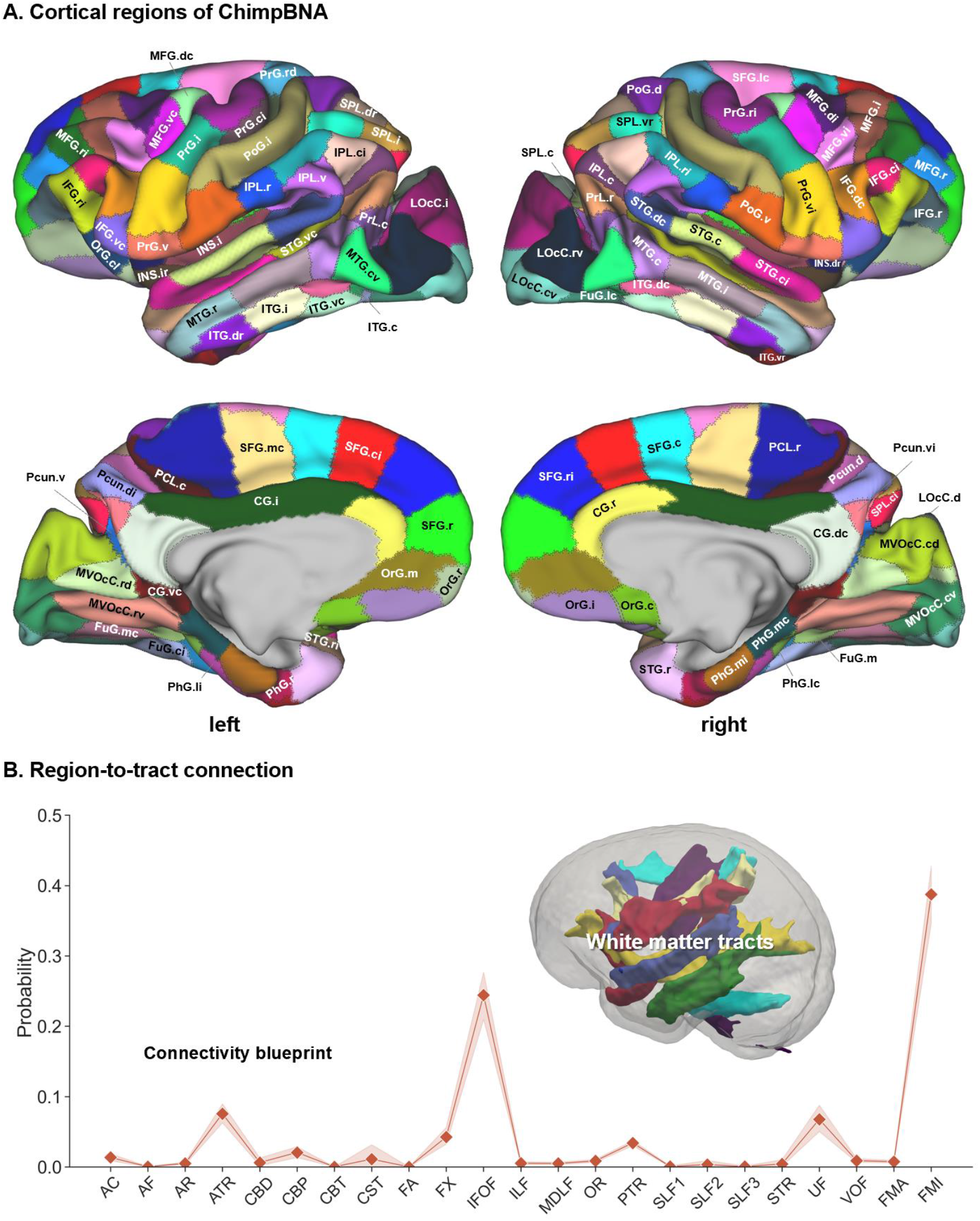
The Chimpanzee Brainnetome Atlas and connections of the chimpanzee brain. **(A)** Cortical regions of the Chimpanzee Brainnetome Atlas. 100 cortical subregions were identified per hemisphere using anatomical connectivity profiles. **(B)** Region-to-tract connections of the chimpanzee brain. White matter tracts were reconstructed following protocols in the previous study ^44^, and the connectivity blueprint of an exemplar region, SFG.r, is shown. SFG.r, superior frontal gyrus, rostral part.

Considering the homology of sulci and gyri between chimpanzees and humans, the definition and naming of initial regions of the chimpanzee brain followed that of the human brain when constructing the Human Brainnetome Atlas (HumanBNA) ^37^, where 20 initial cortical seeds were delineated. Here, we merged one seed, i.e., the posterior superior temporal sulcus (pSTS), into other initial regions in the temporal lobe due to its uncertain definition in chimpanzees, thus obtaining 19 cortical seeds (Table S1). The boundaries between two large gyri were manually edited at the mid-point of the sulcus. Due to the scarcity of chimpanzee brain atlases that could be employed for referencing the labeling schemes, we named the subregions according to their topological positions. The terminology of the parcellation is provided in Table S2. Although detailed whole-brain cytoarchitectonic data from chimpanzee brains are unavailable for alignment, we utilized the cytoarchitectonic maps of three well-described regions for validation ^21,40,46–49^ and found good consistency between histology and connectivity-base parcellation (inferior parietal lobule, Figure S4A; inferior frontal gyrus, Figure S4B; superior temporal gyrus, Figure S4C). The details of the parcellation results for each initial region are shown in Figure S5-S30.

To evaluate the validity of the ChimpBNA, distance-controlled boundary coefficient (DCBC) was used to measure the homogeneity and heterogeneity of the parcellations using structural connectivity ^50^. DCBC is an unbiased criterion that can compare within-parcel to between-parcel similarity considering the spatial autocorrelation of the brain data and the scale of the parcellations. A higher DCBC indicated that the parcellation provided a brain topography with higher within-parcel similarity and higher between-parcel dissimilarity. We compared the DCBC of ChimpBNA using the structural connectivity of each vertex/voxel with other parcellations of chimpanzees, including Bailey and Bonin’s parcellation (BB38) ^40^, and Davi130 parcellation ^12^. The results demonstrated that ChimpBNA achieved the best performance for the delineation of the chimpanzee brain based on structural connectivity (paired t-test, all *p* < .001, Bonferroni corrected; Figure S31). To reveal the connectivity patterns of the subregions of ChimpBNA, we first obtained a region-to-region connectivity matrix (Figure S32). Each term represented the structural connectivity between subregions and was the count of the connected streamlines of probabilistic tractography. More details of the intra- and inter-hemispheric connections between regions are shown in Figure S32A-C. Taking the left SFG.r as an example, this area had the majority of its connections with frontal subregions, including the SFG.ri, MFG.r, IFG.r, OrG.m, CG.r, and other subcortical regions in the ipsilateral hemisphere and with the SPL.c, INS.rd, INS.cd, and CG.vc in the contralateral hemisphere through the corpus callosum (Figure S32D).

We further assessed the structural hemispheric asymmetry of the chimpanzee brain for each ChimpBNA subregion. The majority of subregions with greater gray matter volume in the left hemisphere after adjusting for Bonferroni correction were located in the rostral inferior parietal lobule (IPL), insula, rostral fusiform gyrus (FuG), and posterior middle frontal gyrus (MFG), while gray matter volumes in the right were caudal IPL, anterior MFG, middle temporal gyrus (MTG), and part of lateral occipital gyrus (Figure S33A). Leftward surface area asymmetry patterns were found in rostral IPL, anterior temporal lobe, and medial occipital gyrus, while rightward asymmetries were found in posterior IPL, anterior cingulate, and MTG (Figure S33B). The hemispheric asymmetric results showed consistent patterns with previous studies ^12,32,33^.

We then reconstructed 45 homologous white matter tracts of the chimpanzee brain following the protocols in a previous study ^44^ and performed probabilistic tractography from each vertex of the white/gray matter surface to the whole brain. The regional connectivity blueprint was then derived by multiplying the unwrapped white matter tracts matrix by the whole-brain connectivity matrix ^45^, with the middle cerebellar peduncle (MCP) disregarded because of its absence of projection to the cortex. The rows showed the region-to-tract connectivity pattern of each subregion. Taking the left SFG.r as an example again, the subregion was mainly connected with the forceps minor (FMI), left inferior fronto-occipital fascicle (IFOF), and anterior thalamic radiations (ATR) (Figure 2B).

### Connectivity Divergence between Species

Leveraging connectivity blueprints built for chimpanzees and humans, we explored the connectivity divergence between these two species. A modified dissimilarity measure used to quantify the difference between two probability distributions, the symmetric Kullback-Leibler (KL) divergence ^45^, was calculated to measure the dissimilarity of the regional connectivity blueprints of the two species between each subregion in the ChimpBNA and the HumanBNA. The minimum divergence of each subregion of the HumanBNA was denoted as the connectivity divergence of that region, resulting in a connectivity divergence map (Figure 3A, S34). Higher values in the divergence map indicated that the region in humans had a connectivity pattern that was more dissimilar to the regions in chimpanzees, i.e., might not be represented in this close phylogenetic relative ^45^. One should note that cladistic inferences about the direction of evolutionary change in trait values cannot be made because the brain connectivity of the last common ancestor is not known. The minimum divergence of each subregion of the ChimpBNA was also calculated (Figure S35).

**Figure 3.**
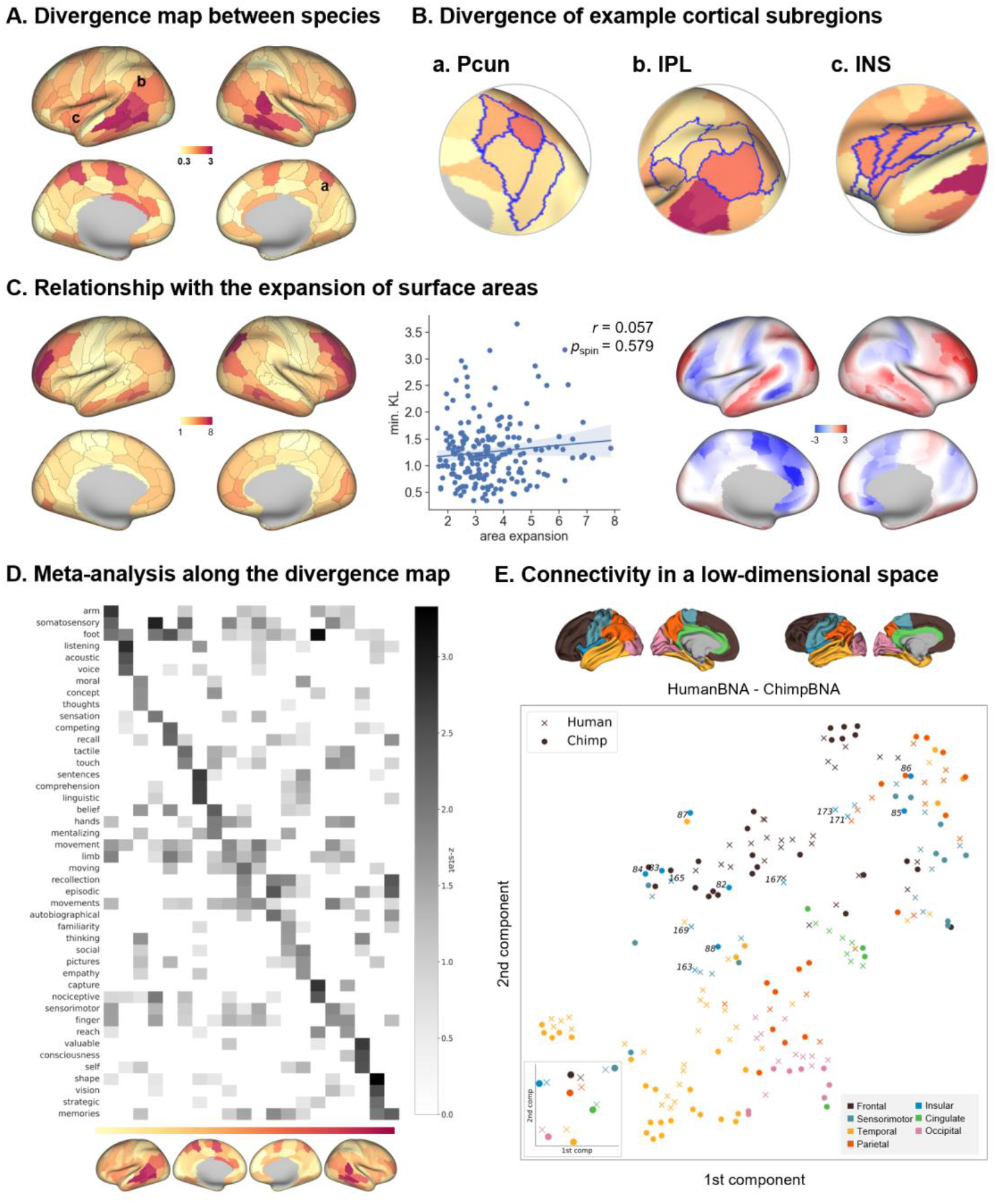
Connectivity divergence between species. **(A)** Connectivity blueprints were used to calculate the KL divergence between species to determine the extent of dissimilarity between their connectivity profiles. Higher values in the divergence map indicated that the region in humans had a connectivity pattern that was more dissimilar to the regions in chimpanzees. **(B)** Connectivity divergence of several example ROIs, i.e., Pcun, IPL, and INS, were investigated at the subregion level. **(C)** The connectivity divergence map showed a very low correlation with the map of cortical expansion between chimpanzees and humans ^13^. The right panel showed the weighted local correlation between the area expansion map and the connectivity divergence map. Regions with high correlation coefficients indicate marked both cortical expansion and connectivity differences between chimpanzees and humans. **(D)** The divergence map was input for functional decoding. NeuroSynth terms with the highest three z-scores for each binarized mask of the divergence map were visualized. **(E)** Subregions of chimpanzees and human left brains were projected into a low-dimensional space using their connectivity profile, with each color representing a cortical system. The figure inset indicates the center of each system. Pcun, precuneus; IPL, inferior parietal lobule; INS, insular cortex.

As shown in Figure 3A, regions with the most different connectivity patterns between species were located at the middle and posterior temporal lobe, especially in the anterior superior temporal sulcus (aSTS) and both the rostral and caudal portion of the posterior superior temporal sulcus (rpSTS and cpSTS), the caudal part of the IPL (corresponding to A39rv or PGa ^51^), the anterior part of the precuneus (Pcun), the insula, and the inferior frontal gyrus (IFG), especially dorsal area 44 (A44d). In contrast, the occipital and sensorimotor cortices showed lower divergence. Notably, we found that the connectivity divergence map showed a very low correlation with the map of cortical expansion (*r* = 0.057, *p*_spin_ = .579; Figure 3C, S36) ^13^, although several regions in the prefrontal cortex showed both morphological and connectional divergence (Figure 3C, right). This indicated that the connectional changes reflect a unique aspect of brain reorganization.

We next grouped the cortical subregions into seven intrinsic functional networks, i.e., the visual (VIS), somatomotor (SMN), limbic (LN), dorsal-attention (DAN), ventral-attention (VAN), frontoparietal (FPN), and default mode network (DMN) (Figure S37B, left) ^52^. Higher-order cognitive networks displayed greater divergence than the VIS/SMN (Mann-Whitney U test, *p* = .0045; Figure S37B, right), with the FPN having the greatest divergence.

Taking the precuneus as an example, the anterior (A5m) and the dorsal-middle (A7m) subregions showed greater connectivity divergence than the ventral-middle (A31) and posterior (dmPOS) subregions, highlighting the connectivity dissimilarity between chimpanzees and humans in this heterogeneous region (Figure 3B, a). Different connectional patterns of the superior longitudinal fasciculus I (SLF1), IFOF, and acoustic radiation (AR) contributed to the divergence between chimpanzees and humans (Figure S38A, B). As for the IPL, the anterior (A40rv) and posterior (A39rv) subregions showed greater divergence than the other subregions in the IPL (Figure 3B, b; Figure S38C, D), and in the insular cortex, the anterior subregions presented a greater difference than the posterior subregions (Figure 3B, c; Figure S38E).

We used NeuroSynth to perform a meta-analysis to investigate the relationship between the divergence maps and various cognitive functions examined in human subjects. This analysis showed that regions with high divergence between humans and chimpanzees were characterized by higher-order functions, such as "memories, " "strategic, " and "shape." In contrast, the lower end of the divergence map was related to more basic sensory and motor processing, including "arm," "somatosensory," and "foot" (Figure 3D).

We further compared atlases of chimpanzees and humans using connectivity blueprints as homologous features. For visualization, we projected regional connectivity blueprints of the two species to a low-dimensional space using t-SNE and visualized each region in a 2-dimensional space (Figure 3E). Regions with similar connectivity profiles were grouped together in the resulting space, and regions belonging to one of the cortical systems (frontal, sensorimotor, temporal, parietal, insular, cingulate, and occipital) were labeled with distinct colors. The sensorimotor, cingulate, and occipital regions tended to form a group, but the other regions were more scattered, especially the insular subregions, which were scattered among different cortical systems (Figure 3E). The median coordinates of each cortical system were also calculated (Figure 3E, inset). The primary cortex, including the sensorimotor and occipital regions, was closer between chimpanzees and humans, while the frontal and parietal regions were farther apart, with the largest dissimilarity in the temporal regions.

### Whole-brain level Connectional Lateralization

To investigate the connectional lateralization of subregions of the chimpanzee brain, we split the connectivity blueprint into two hemispheres, *CB_L_* and *CB_R_*, which only contained the respective left- or right-hemispheric tracts and the commissural tracts (21 unilateral tracts and 2 commissural tracts). We calculated the KL divergence between the homotopic subregions of the hemispheres of the chimpanzees, indicating the inter-hemispheric differences in connectivity patterns. As shown in top left panel of Figure 4A, the regions with highly asymmetric connectivity patterns included the posterior temporal lobe, IPL, medial occipital cortex, and MFG. Inter-hemispheric differences in human brain regions were also calculated the same as in the chimpanzee data, and the patterns were largely consistent with previous studies (Figure 4A, bottom right) ^53^.

**Figure 4.**
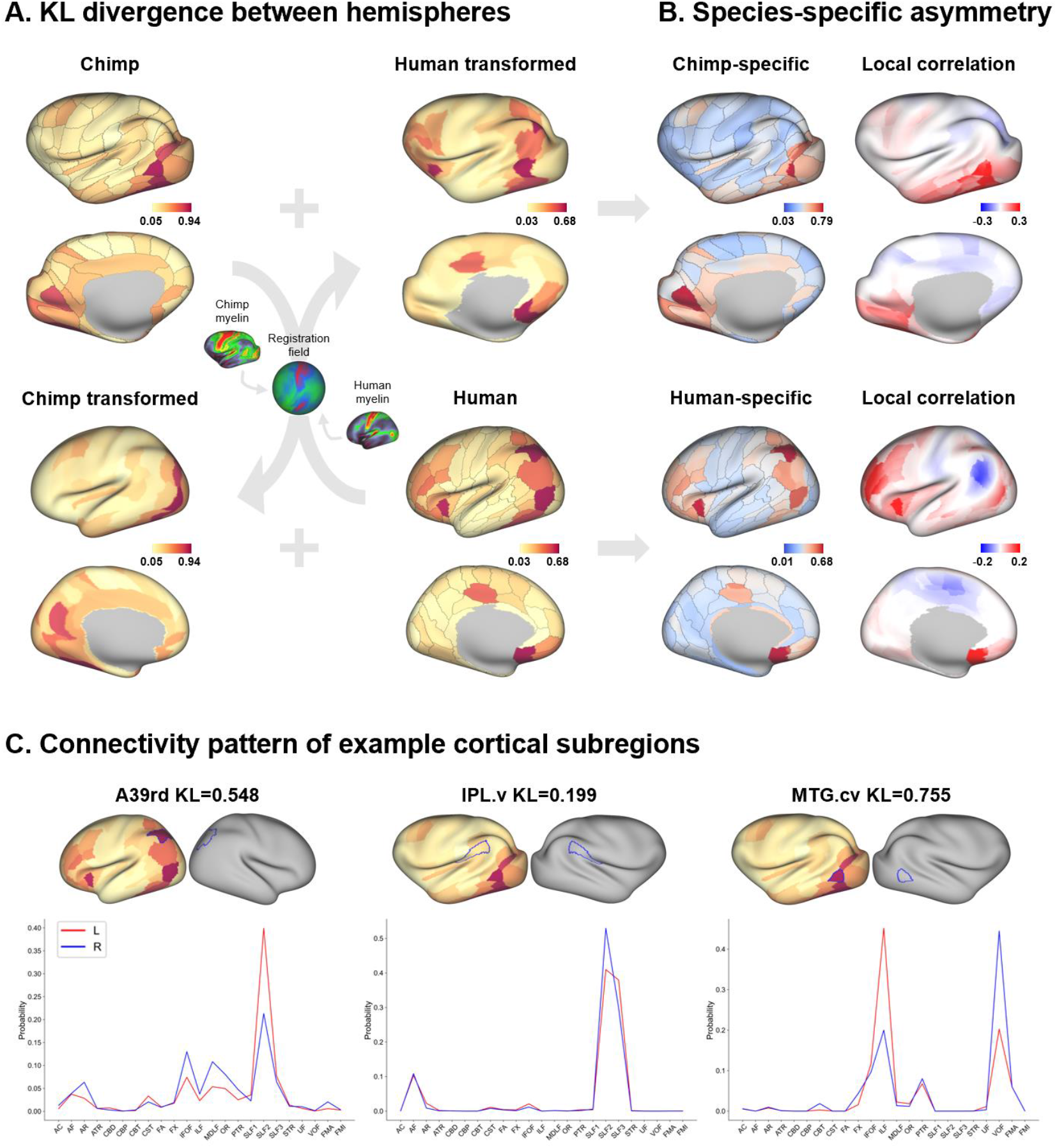
Species-specific whole-brain level connectional lateralization. **(A)** Connectivity blueprints were used to calculate the KL divergence between homotopic subregions between hemispheres. The divergence map of the connectional lateralization of the chimpanzees and humans was aligned and compared. **(B)** Species-specific asymmetric connectivity patterns were calculated, and the weighted local correlation with the connectivity divergence map is shown, where a higher value indicates both marked connectivity differences between species and asymmetry between hemispheres. **(C)** Connection probability of tracts of several example ROIs. In A39rd in humans, the inter-hemisphere differences were driven by IFOF, MdLF, and SLF2, while in IPL.v in chimpanzees, the asymmetry was mainly driven by SLF2. MTG.cv in the chimpanzee brain, in which the KL divergence was greater, showed differences in their tract connections, mostly driven separately by the ILF and VOF. A39rd, rostrodorsal area 39; IPL.v, inferior parietal lobule, ventral part; MTG.cv, middle temporal gyrus, caudoventral part; IFOF, inferior fronto-occipital fasciculus; MdLF, middle longitudinal fasciculus; SLF2, superior longitudinal fascicle II; ILF, inferior longitudinal fasciculus; VOF, vertical occipital fasciculus.

Since the data for the two species were not in common space and could not be compared directly, we used an alignment technique based on myelin data to transform the asymmetric connectivity pattern between chimpanzees and humans ^14,54^. The cross-species alignment was based on a multimodal surface matching algorithm (MSM) ^55^, using the distinction of areas in myelin maps as anchor points, and derived a cortical registration between species. This allowed us to compare species-shared and species-specific regions with asymmetric connectivity patterns in a common space.

In the common human space, we focused on the human-specific regions that had asymmetric connectivity patterns by calculating a variant of the exclusive *OR* maps between the chimpanzee and human brains (Figure 4B) ^56^. As shown in the bottom left panel of Figure 4B, the human maps showed unique regions with asymmetric connectivity in the dorsal part of the IPL, anterior part of the insular cortex, middle cingulate cortex (MCC), posterior part of the orbitofrontal cortex (OFC), and most of the lateral prefrontal cortex (PFC). Weighted local correlation maps between the human-specific asymmetry pattern and the connectivity divergence map provide a visualization of the reorganization of the brain connectivity between the species. Regions with high values in the human brain indicate marked connectivity differences between chimpanzees and asymmetry between hemispheres. The posterior inferior parietal lobule, temporal lobe, posterior orbitofrontal cortex, and dorsolateral prefrontal cortex showed the greatest differences (Figure 4B, bottom right).

Similarly, we found the chimpanzees showed unique connectional asymmetries in the several posterior temporal regions and medial occipital cortex (Figure 4B, top left), and showed marked connectional changes in the caudoventral part of the temporal lobe (Figure 4B, top right), which could be considered chimpanzee-unique connectional features distinct from human.

We next investigated the tract contribution to the asymmetric pattern for several example subregions. For the rostrodorsal area 39 (A39rd) in humans, which showed human-specific asymmetric connectivity pattern, the asymmetric tract IFOF, middle longitudinal fasciculus (MdLF) and superior longitudinal fasciculus II (SLF2) showed their contribution (Figure 4C, left), while for the IPL.v, which also showed an inter-hemispheric difference in chimpanzee, the asymmetry was mainly driven by the SLF2 (Figure 4C, middle). This is consistent with the finding that the C2 subregion in the inferior parietal lobule of chimpanzees showed significant rightward asymmetric connections with the SLF2 ^33^. The SLF2 also contributed to the asymmetry observed in MFG (Figure S39A). For the highly asymmetric MTG.cv in chimpanzees, the inferior longitudinal fascicle (ILF) and vertical occipital fascicle (VOF) showed distinct lateralized connections (Figure 4C, right). The connectional asymmetries in MVOcC.rd and MVOcC.cv were driven by the significant symmetric connections with optic radiation (Figure S39B).

### Gene Associations with Connectivity Divergence between Species

We next investigated the association between the connectivity divergence map and gene expressions using the Allen Human Brain Atlas (AHBA) ^57^. Note that the gene analysis was limited to left hemisphere because few samples were obtained from the right hemisphere in AHBA. We used partial least squares regression (PLSR) on the AHBA data and generated the first component (PLS1 score). The PLS1 score significantly correlated with the divergence map after a 10,000 times spin test (*r* = 0.39, *p*_spin_ < .011; Figure 5A). 1939 genes with a *Z*-score greater than 3 were filtered using 10,000 times bootstrapping, including *EFCAB1*, *NUDT11*, *C2CD4C*, and *SYT17* which showed higher *Z*-score (Table S3). These genes were enriched in excitatory neurons that showed the highest level of significance, followed by astrocytes (Figure 5B). Specifically, the L6 and L2-3 intratelencephalic excitatory neurons were the most strongly enriched. The genes with a *Z*-score higher than 3 were related to neuronal formation, neuron projection, and synapses (Figure 5C).

**Figure 5.**
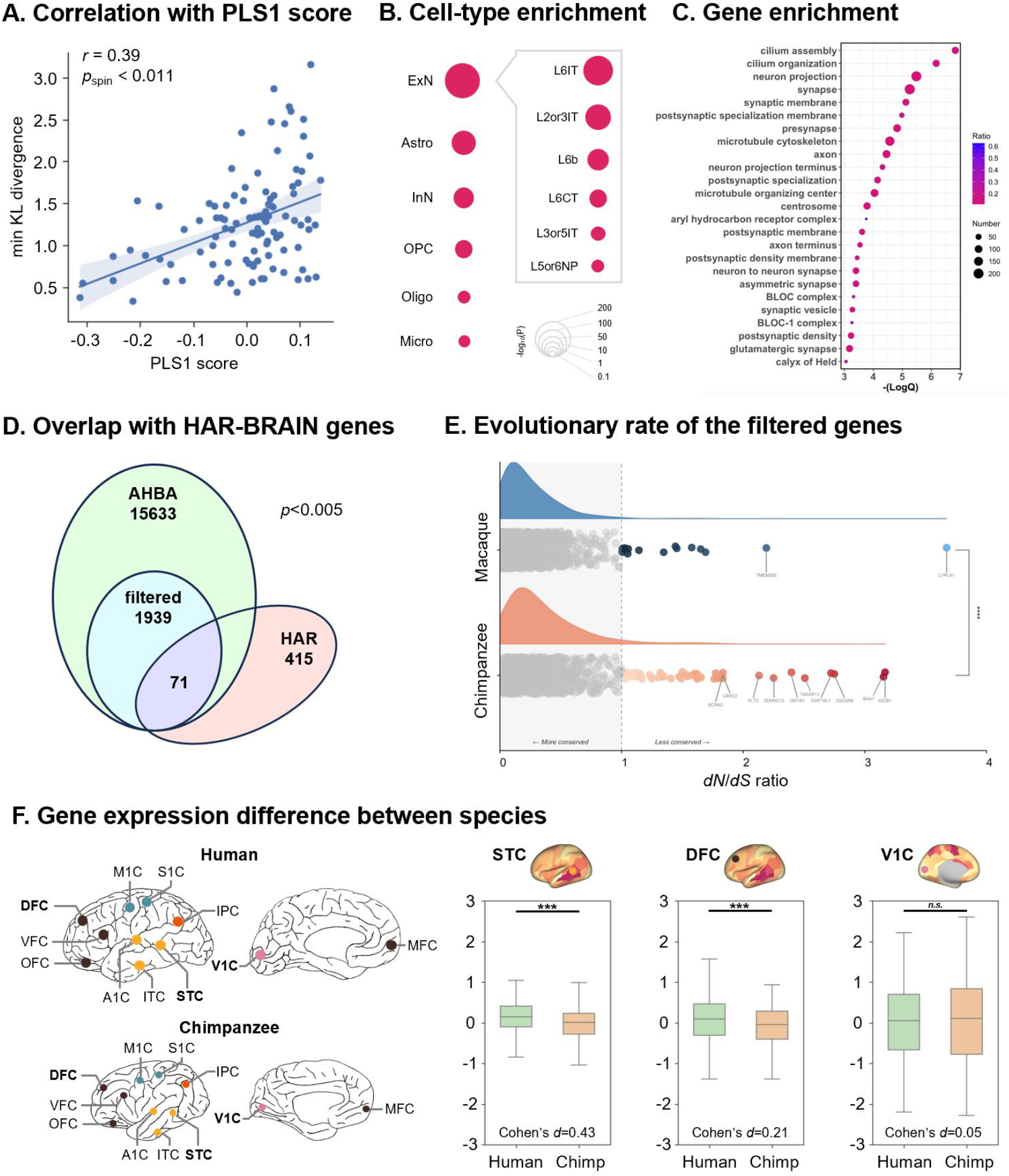
Gene association with connectivity divergence between species. **(A)** The divergence map shows a significant correlation with the PLS1 score of genes with the AHBA dataset (left brain). **(B)** Cell-type enrichment analysis of genes identified using bootstrapping with the most positive or negative weights (|*Z*| > 3) in PLSR. **(C)** These genes were used in an enrichment analysis and found to be associated with neuronal projection and synapse formation processes. **(D)** 71 genes overlapped with the HAR-BRAIN genes (*p* < .005). **(E)** 56 genes had a *dN*/*dS* ratio > 1 (greater than 1 means less conserved in chimpanzees), but only 14 genes in the macaque against the human genome (Welch’s t-test *p* < .0001). **(F)** Differences in these filtered genes were compared using human and chimpanzee data from the PsychENCODE database. 1473 of 1939 genes that overlapped in the database were used for analysis. Three regions of interest exhibiting distinct connectivity divergence were considered, and a paired t-test was utilized to assess differences between the two species. Significant differences were found in first two regions, and the effect size was more significant in the STC and DFC than in V1C. STC, superior temporal cortex; DFC, dorsolateral frontal cortex; V1C, primary visual cortex. *** indicates *p* < .001.

In addition, 71 of these genes significantly overlapped with human-accelerated genes related to brain processes (HAR-BRAIN genes ^13^) (*p* < .005; Figure 5D, Table S4). We also quantified the evolutionary rate, i.e., *dN*/*dS* ^58^, of these genes to examine whether these genes experienced positive selection (*dN*/*dS* > 1), neutral selection (*dN*/*dS* = 1), or negative selection (*dN*/*dS* < 1) since the last common ancestor of humans and chimpanzees. Macaque (*Macaca mulatta*) data was also utilized for comparison with chimpanzee-human results. Fifty-six of 1939 genes were found to have *dN*/*dS* > 1 in the chimpanzee-human clade, whereas only 14 genes with positive selection were found for the macaque data (Welch’s t-test, *p* < .0001; Figure 5E, Table S5).

We further examined whether these filtered genes show differential expression between humans and chimpanzees. We used comparative gene expression data from the PsychENCODE database ^26^, which contains the expression levels of 16463 genes at 16 homologous brain locations in humans (*n* = 6), chimpanzees (*n* = 5), and macaques (*n* = 5). Of the 1939 genes with *Z*-scores above 3, 1473 overlapped with the PsychENCODE data and were used for further study (Table S6). Gene expression data were normalized across the cortical areas to obtain *Z* scores and averaged to a group-level gene expression for humans and chimpanzees. We selected three regions of interest that showed distinct connectivity divergence in our study, i.e., the superior temporal cortex (STC), dorsolateral frontal cortex (DFC), and primary visual cortex (V1C). We first assessed the two species’ differences using a paired t-test for each region. We found significant differences in first two regions (STC: *t* = 16.26, *p* < .001, Bonferroni corrected; DFC: *t* = 8.16, *p* < .001, Bonferroni corrected; V1C: *t* = -1.88, *p* = .0599) and more significant effect size in the STC and DFC than in the V1C (STC: Cohen’s *d* = 0.42; DFC: Cohen’s *d* = 0.21; V1C: Cohen’s *d* = 0.05). The differentially-expressed genes between species for each region were called at a FDR of 0.01 (122 genes in STC, 92 genes in DFC, 118 genes in V1C; Figure S40) and were not attributable to variations in the ratio of major cell types between species (Figure S41). This indicates a potential association between gene expression differences and connectivity divergence between species.

## Discussion

This study systematically compared the connectivity profiles of chimpanzees and humans, revealing divergent connectivity patterns in wiring space across species, the potential genes involved, and the asymmetric connectivity pattern at the whole-brain level. To enable cross-species comparisons, we first developed a fine-grained atlas of the chimpanzee brain, ChimpBNA, based on anatomical connectivity information and following our previous framework developed for the human brain. We subdivided the chimpanzee brain into 122 subregions (100 cortical and 22 subcortical regions) per hemisphere and mapped the structural connectivity pattern for each subregion. We used these connectivity blueprints to examine connectivity divergence between chimpanzees and humans. Specifically, we found that the two species showed profound dissimilarities in the posterior temporal lobe, IPL, Pcun, insula, and IFG. We next explored the lateralization of the connectivity patterns for each species and showed human-unique asymmetries in the insula, OFC, and lateral PFC, and chimpanzee-specific asymmetries in the caudoventral temporal regions and medial occipital. The genes filtered according to connectivity divergence were especially enriched in cells involved in the construction of cortical projection circuits and the formation of synapses. The genes showed different evolutionary rates and expression patterns between the species, suggesting a potential link between gene expression and connectivity.

### Fine-grained Chimpanzee Brain Parcellation using Anatomical Connectivity

Atlases are essential for investigating the structural and functional characteristics of the brain by providing a map in a common space and allowing comparison of results across studies ^59^. A series of comparable brain atlases with a uniform parcellation scheme from different species could facilitate efficient comparative studies and provide vital clues about how the human brain evolved ^60,61^. However, mapping non-human primate brains is incomplete and relatively preliminary, with previous atlases having been constructed using different modalities and at different scales. This hinders the translation of results across species and the understanding of human brain structure and function ^35^. Although a few atlases have been developed for chimpanzee brains ^12,21,40,41^, a fine-grained parcellation utilizing a uniform scheme comparable to humans has been lacking. To address this deficiency, we constructed the ChimpBNA using anatomical connectivity profiles, which will improve our understanding of the anatomical brain organization of chimpanzees and enable comparative analysis across species.

The earliest chimpanzee whole brain atlas dates back to the 1950s when Bailey and his colleagues delineated cortical regions based on histological sections ^40^. Although later mapped to MRI slices, the information available is limited due to its coarseness ^21,41^. A more recent parcellation, Davi130 ^12^, was manually segmented based on macroanatomical landmarks but lacks connectivity information on its subregions. While acknowledging that there is no consensus as to which type of information reflects the true anatomical organization of the brain ^62,63^, the ChimpBNA provides a whole-brain parcellation of the chimpanzee brain into robust and biologically plausible subregions, together with detailed characterizations of structural connectivity patterns for each area. Anatomical connections confirmed the boundaries previously detected by cytoarchitecture and identified several subdivisions not previously reported in brain mapping of human and other non-human primates ^37,64^. For example, the IPL contained 5 subregions in our atlas (Figure S4A, Figure S17) and showed consistent rostral-caudal topological pattern across various parcellation schemes, except for a ventral subregion located in the parietal operculum, which has been reported both by anatomical connection-based and cytoarchitectonic parcellation ^33,46,65^. For another example, the chimpanzee’s IFG has received extensive attention because of its potential association with the evolution of communication and language ^66,67^. Previous studies demonstrated that a homolog of Broca’s area is present in chimpanzee brains, but interindividual variability in the boundaries of area 44 and area 45 does not align with morphological sulcal anatomy ^68^. Here, we segregated the IFG into 6 more refined subregions: two dorsal and a ventral portion of putative area 44, the rostral and caudal parts of putative area 45, and one most rostral cluster approaching the frontal pole (Figure S4B, Figure S7), providing a potential reference for further analysis of this area. Meanwhile, another language-related homolog in the chimpanzee brain, the area Tpt, a component of Wernicke’s area ^48^, was also present in the posterior STG in the ChimpBNA (Figure S4C, Figure S11).

### Connectivity Divergence between Species

Comparing evolutionary brain changes is fundamental for understanding anatomical and functional similarities and differences across species ^69^. Although the expansion of brain areas and resulting reorganization between chimpanzees and humans has been investigated ^13^, the divergence in connectivity patterns between humans and their closest primate relatives in the evolutionary tree has yet to be comprehensively analyzed. The connectivity blueprints framework provides a common space to enable comparisons of cortical organization and white matter connectivity across species, quantitatively identifying not only common principles but also unique specializations throughout evolution ^45^.

The current study presents a connectivity divergence map showing cortical regions such as the middle and posterior temporal lobe, inferior parietal, precuneus, insular, and inferior frontal gyrus with particularly higher connectivity divergence between humans and chimpanzees. Functional decoding analysis using NeuroSynth data indicated that these regions were related to "memories," "shape," "strategic," and "self" ^70–72^ when assessed in humans. That said, it is important to recognize that this analysis is derived from human data alone. The functional role that these pathways have in chimpanzee cognition has not been studied, so it is difficult to make direct comparisons between species. Further, humans and chimpanzees are different in many other anatomical and physiological features (i.e., bipedalism, vocal control, etc.) that might be related to evolutionary changes in connectivity that are not captured in the Neurosynth meta-analytic approach we adopted in this paper. These limitations aside, given the evidence that evolutionary changes in anatomical connections may contribute to functional differences in neural systems across species ^16,17,20,21^, the overall results point to divergences in neuroanatomy that may potentially support unique species-specific cognitive and motor specializations ^73^.

The precuneus is a heterogeneous region that is implicated in complex cognitive functions. The anterior (A5m) and middle (A7m) subregions, which are involved in sensorimotor and cognitive processes, had a greater divergence than the posterior subregion (dmPOS), which is associated with visual processing ^74^. Chimpanzees generally have stronger and more flexible limbs to maneuvur in their environment, which could cause the difference in the anterior subregions, while humans outperform apes in various cognitive tasks ^75^.

As for the lateral parietal cortex, previous studies have shown that the parietal operculum of chimpanzees exhibited asymmetry and was involved with handedness in the use of tools ^76^. Although non-human primates share many tool capabilities, humans uniquely show greater manual dexterity and conceptual knowledge of tool use ^70^. Considerable divergence was found in the tool processing network nodes ^70,77^, the inferior frontal gyrus, and the most rostroventral inferior parietal lobule (A40rv). These two regions were connected by the IFOF and SLF3, which have been linked with the human tool-use circuit ^78–80^ and showed a different connectivity pattern between chimpanzees and humans. While the rostroventral group of areas of the IPL deals with tool use and sound, the middle and caudal IPL are implicated in nonspatial attention processes and semantic processing, respectively ^51^. This is reflected by greater divergence in the A39rv (PGa) than in other subregions of the IPL.

Of the cortical regions that showed the greatest connectivity divergence, the temporal lobe merits special attention due to the significant changes that have been described in this region over primate evolution ^20,81–84^. Although the temporal cortex does not show as much cortical expansion as the frontal cortex, there was a significant connectivity difference between chimpanzees and humans in the temporal cortex. The posterior temporal lobe is involved with several human-unique cognitive abilities, including language comprehension. The arcuate fasciculus, which is known to be associated with language processing ^85^, has been described to extend further anteriorly and inferiorly in the temporal lobe in humans than in chimpanzees, resulting in the alternation of connectivity patterns in this area, which might support the evolutionary specialization of this area ^14,44^. These findings suggest that these regions may have been expanded and reorganized in human brains and resulted in the emergence of more complex linguistic abilities, notably speech processing.

We also compared the atlases of the two species by projecting the connectivity blueprints into a low-dimensional space, linking the human and chimpanzee brains into a common connectivity space. Subregions from the same cortical systems tended to group together. However, some regions belonging to a system, e.g., the insular subregions, were more scattered. The posterior insular cortex connected reciprocally with the secondary somatosensory cortex, while the anterior insular cortex interconnected with regions in the temporal and ventral frontal cortex ^86^, indicating posterior-anterior trends of connectivity divergence in the insula. Subregions of the insular cortex were scattered near the frontal, temporal, and sensorimotor systems in the low-dimensional space, indicating the widely different functions related to the insular cortex, from multisensory information processing to the engagement of social emotions and flexible behaviors ^87^. In addition, the primary sensory and motor cortical areas, which showed less connectivity divergence, were closer between chimpanzees and humans than the frontal and parietal regions, which showed a greater distance, with the greatest dissimilarity in the temporal regions. Our results suggest that the additional evolutionary adaptations for the connectivity of the temporal cortex may play a unique role in human specialization ^20,84^.

### Asymmetric Connectivity Differences between Chimpanzees and Humans

Recent neuroimaging studies have shown that hemispheric asymmetry in structural characteristics ^32^, connectivity patterns ^33,34^, and functional activity ^88,89^ are not confined to humans but are actually widespread among non-human primates. Here, we built whole-brain connectivity blueprints, which represent the anatomical connectivity pattern of each subregion using homologous white matter tracts across species, to investigate the inter-hemispheric difference for each species. We found that chimpanzees showed highly asymmetric connectivity patterns in the posterior lateral temporal lobe, inferior parietal, medial occipital cortex, and middle frontal gyrus. In contrast, humans showed highly asymmetric connectivity patterns in the posterior temporal lobe, the superior and inferior parietal cortex, the anterior insular cortex, the posterior orbitofrontal cortex, and most of the lateral prefrontal cortex.

Previous studies reported that chimpanzees showed few leftward asymmetric connections between the IPL and the temporal cortex ^33^, including the planum temporale, anterior superior temporal gyrus, and anterior superior temporal sulcus, all of which have been implicated in human language comprehension ^48^ as well as semantic and phonologic processing ^20,90^. Tract analysis showed that the IPL.ri displayed significant rightward asymmetric connections with the SLF2, which roughly corresponded to the conclusion of the C2 cluster of the IPL in a recent study ^33^ and was implicated in auditory perception, which is one aspect of language processing ^91^. The SLF2 connects the angular gyrus and middle frontal gyrus of the chimpanzee brain ^44^. The posterior portion of the middle frontal gyrus, including the MFG.vi, MFG.dc, and MFG.vc, which likely correspond to PB and PC in Bailey et al. ^40^, also showed asymmetric connections in the SLF2 (Figure S39A). These regions were reported to be more activated by observing an action than by producing grasping both transitively and intransitively ^92^. Asymmetric connectivity in the MTG.cv was supported by the ILF, whose left volumetric asymmetry below the temporal cortex might increase the left connectivity more than the right ^93^. The MVOcC.rd and MVOcC.cv, which were situated in the ventral part of the cuneus cortex and the calcarine sulcus, showed significant asymmetric connections with optic radiation (Figure S39B). The asymmetric connection of the subregions might be based on the asymmetric white matter volume underlying the occipital gyrus^93^. We note here that this study consisted of 37 right-handed and 3 left-handed human subjects, while no attempt was made to match chimpanzees on the basis of handedness. Thus, one might argue differences in the distribution of right- and left-handed humans and chimpanzees might explain some of the findings on divergence in asymmetry between the species. Recent studies in humans have reported that individual differences in handedness are weakly associated with structural asymmetries ^94^. Therefore, though we cannot rule out this explanation, we do not believe that this factor has a significant impact on the divergence patterns reported here.

Given the diversity of the brain in many aspects across species, a common space is required to address these considerations. Based on myelin data, we aligned the chimpanzee brain with the human brain into a common space and explored species-specific regions with asymmetric connectivity patterns. Human uniqueness in asymmetric connectivity was found in the dorsal part of the IPL, anterior insular cortex, MCC, posterior OFC, and most of the lateral PFC. In humans, these regions are involved in tool use ^95^, empathy ^96^, planning ^97^, and abstract reasoning ^98^. Though chimpanzees have been reported to exhibit each of these abilities, the available evidence suggests that their aptitudes in these domains of function are more limited compared to humans ^99^. The posterior inferior parietal lobule, the temporal lobe, and the posterior orbitofrontal cortex all showed distinct asymmetry between hemispheres and connectivity divergence between species, suggesting a possible starting point for explaining the functions executed by these regions, with the emergence of hemispheric specializations in human evolution. However, it is important to note that the observed distinct species-specific patterns based on this comparative analysis cannot resolve the direction of evolutionary changes on the human or chimpanzee lineage due to uncertainly in determining brain connectivity of their last common ancestor; therefore, caution should be exercised when interpreting these results in a phylogenetic context.

### Genetic Factors Associated with Connectivity Divergence between Species

The connectivity divergence and its underlying genetic associations were further investigated. The filtered genes identified by PLSR using the AHBA dataset are implicated in synapse and axon-related processes and neuronal projections, which are at the basis of macroscale anatomical connections ^25,100^. Several genes, including *EFCAB1*, *C2CD4C,* and *SYT17*, were predicted to enable calcium ion binding ^101,102^, which plays an important role in signal transduction and other cellular processes. *SYT17* is also involved in positively regulating dendrite extension and enabling syntaxin binding activity ^103^. *TMSB10* is related to actin monomer binding activity, thus regulating cell migration ^104–106^. Other genes that had a greater weight were related to metabolism or regulation of transcription factor activity.

The filtered genes were enriched in excitatory neurons in the upper (L2/3) and deeper (L6) layers as well as being enriched in astrocytes. The excitatory pyramidal neurons contribute to the construction of major cortical projection circuits ^107^, and astrocytes are involved in regulating synaptic transmission and plasticity ^108^. Our findings suggest that potential interactions between excitatory neurons from different cortical layers and astrocytes contribute to divergent connectivity between species at the cellular level.

From an evolutionary perspective, the genes shows signatures of positive selection (i.e., *dN*/*dS* > 1) in the chimpanzee-human phylogenetic group (i.e., the Hominini clade) compared with macaques and markedly overlapped with the HAR-BRAIN genes, which are enriched in human-evolved elements that are associated with brain development, cortical expansion, and formation of connections ^13,109^. Chimpanzees share a more recent common ancestor with humans than macaques, so they display more similar genetic changes due to their phylogenetic proximity. They are more likely to have shared genetic adaptions and positive selection events, especially those related to hominoid traits ^110,111^. These genes were expressed unequally in the cortex, with more significant differences in regions with a greater connectivity divergence. This suggests that the genes behind the connectivity divergence played an essential role in modern human evolution and may reflect increased cognitive specialization across species. However, we note that the difference in detected gene transcript abundance levels may be due to differences in cellular composition between the two species. With the development of single-cell RNA sequencing technology applied to a greater range of brain regions across species ^112,113^, these confounding effects can be distinguished in future studies, and more accurate biological inferences could be made.

### Methodological Considerations

This work has some technical and methodological limitations to acknowledge. The first concern is the false positives produced by tractography ^114^. However, diffusion MRI tractography has been irreplaceable for *in vivo* and non-invasively investigating the organization of humans and other non-human primates, such as chimpanzees. A recent study mapped both deep and superficial white matter in the chimpanzee brain, providing an important understanding of this species ^115^. Together with *ex vivo* neuroanatomical data, the joint analysis may mitigate these technical issues and provide even more insights ^24,116^. Second, given the different brain sizes across species, the incidence of potentially more acute curvatures in some tracts should also be considered. Because of the relatively low-resolution diffusion images of the chimpanzee brain, only large subcortical nuclei were parcellated based on anatomical connectivity. Other smaller nuclei and the cerebellum were not considered in this study. Recent studies have analyzed these non-neocortical areas ^117,118^, which deserve greater attention. Finally, regarding parcellation reliability, it should be acknowledged that the *a priori* macro-anatomical boundaries may not be related to the actual differentiation of cortical areas based on pure connectivity profiles ^37^. Future work needs to quantitatively examine the correspondence between the ChimpBNA and microstructural parcellations in a greater range of brain regions. However, it is difficult to define the number of subregions when parcellating starting from the whole cortex, but the sulci were reported to be reliable determinants of the anatomo-functional organization of brains ^119,120^. Combined with existing primate parcellations ^42,43^, ChimpBNA enables efficient cross-species brain comparisons. However, accurate homologous regional correspondence across species remains lacking despite uniform atlas-building schemes. Establishing more accurate homology mapping across species should be a priority for future work on developing comparable cross-species brain atlases.

## Conclusion

In summary, we constructed the fine-grained ChimpBNA based on anatomical connectivity and performed a systematic comparative analysis of chimpanzee and human connectivity profiles. This revealed divergent patterns across species, associated genetic factors, and whole-brain connectional asymmetry. Distinct connectivity divergences between species were observed in the cortical association areas, including the posterior temporal, inferior parietal, precuneus, insula, and inferior frontal gyri, with related genes showing differential evolution and expression. We also identified shared asymmetric areas constrained to the posterior temporal, inferior parietal, and mid-cingulate cortices, while unique human asymmetry was seen in the insula, orbitofrontal cortex, and lateral prefrontal cortex. The genes filtered according to the connectivity divergence were enriched into cell types related to the construction of cortical projection circuits and involved with the formation of synapses and showed different evolutionary rates and expression patterns across species. Our findings indicate that connectivity divergence in chimpanzees and humans likely reflect functional specialization in evolution and provide key steps toward elucidating the diversity of primate brains.

## Methods

### Data Acquisitions and Preprocessing

#### Human data

Data from 40 healthy human adults (*Homo sapiens*, 17 males) were selected as the same subjects as in the construction of the Human Brainnetome Atlas (HumanBNA) ^37^ from the S1200 subjects release of the Human Connectome Project (HCP) database ^121^ (http://www.humanconnectome.org/). All the scans and data from the individuals included in the study had passed the HCP quality control and assurance standards.

The scanning procedures and acquisition parameters were detailed in previous publications ^122^. In brief, T1w images were acquired with a 3D MPRAGE sequence on a Siemens 3T Skyra scanner equipped with a 32-channel head coil with the following parameters: TR = 2400 ms, TE = 2.14 ms, flip angle = 8°, FOV = 224×320 mm^2^, voxel size = 0.7 mm isotropic. DWI images were acquired using single-shot 2D spin-echo multiband echo planar imaging on a Siemens 3 Tesla Skyra system (TR = 5520 ms, TE = 89.5 ms, flip angle = 78°, FOV = 210×180 mm). These consisted of three shells (b-values = 1000, 2000, and 3000 s/mm^2^), with 90 diffusion directions isotropically distributed among each shell and six b = 0 acquisitions within each shell, with a spatial resolution of 1.25 mm isotropic voxels.

#### Chimpanzee data

Data from 46 adult chimpanzees (*Pan troglodytes*, 18 males) were available from the National Chimpanzee Brain Resource (NCBR, http://www.chimpanzeebrain.org). The data, including T1w and DWI, were acquired using previously described procedures at the Emory National Primate Research Center (ENPRC) on a 3T MRI scanner under propofol anesthesia (10 mg/kg/h) ^123^. All procedures followed protocols approved by ENPRC and the Emory University Institutional Animal Care and Use Committee (IACUC, approval no. YER-2001206). All data were obtained before the 2015 implementation of U.S. Fish and Wildlife Service and National Institutes of Health regulations governing research with chimpanzees. All chimpanzee scans were completed by the end of 2012; no new data were acquired for this study.

T1w images were collected at a 0.7×0.7×1 mm resolution. DWI images were acquired using a single-shot spin-echo echo-planar sequence for 60 diffusion directions (b = 1000 s/mm^2^, TR = 5900 ms; TE = 86 ms; 41 slices; 1.8 mm isotropic resolution). DWI images with phase-encoding directions (left–right) of opposite polarity were acquired to correct susceptibility distortion. Five b = 0 s/mm^2^ images were also acquired with matching imaging parameters for each repeat of a set of DWI images.

#### Data preprocessing

The human T1w structural data had been preprocessed following the HCP’s minimal preprocessing pipeline ^122^, while the chimpanzee and macaque T1w structural data were preprocessed following the HCP-NHP pipelines described in previous studies ^11,124^. In brief, the processing pipeline included imaging alignment to standard volume space using FSL, automatic anatomical surface reconstruction using FreeSurfer ^125^, and registration to a group average surface template space using the multimodal surface matching (MSM) algorithm ^55^. Human volume data were registered to Montreal Neurological Institute (MNI) standard space, and surface data were transformed into surface template space (fs_LR). Chimpanzee volume and surface data were registered to the Yerkes29 chimpanzee template ^11^. All subjects included in this study passed automatic FreeSurfer quality control and visual inspection.

Preprocessing of the diffusion-weighted images was performed in a similar way in the human and chimpanzee using FSL ^126^. FSL’s DTIFIT was used to fit a diffusion tensor model. Following preprocessing, voxel-wise estimates of the fiber orientation distribution were calculated using bedpostx, allowing for three fiber orientations for the human dataset and two fiber orientations for the chimpanzee dataset due to the b-value in the diffusion data.

### Connectivity-based Parcellation

#### Initial seed mask definition

Following the protocols we used in constructing the HumanBNA ^37^, we identified the initial seeds of the chimpanzee brain for the subsequent parcellation. Specifically, we started from the Desikan-Killiany-Tourville (DKT) atlas ^127^, which served as a foundational reference for identifying gyri and sulci that was used as *a priori* information to guide the fine-scale subdivision process. ROIs representing subdivisions of a larger gyrus, as well as distinct anatomical landmarks that the DKT atlas delineates, were combined, and the boundaries between two gyri were manually edited at the mid-point of the sulcus by two authors (W.Y. and C.L.). The resulting 19 cortical seeds were delineated on the Yerkes29 mid-thickness surface. In addition, 7 subcortical regions were identified using FreeSurfer and were included in the analysis. The full names and abbreviations of the initial cortical and subcortical seed masks are listed in Table S1.

#### Connectivity-based parcellation

We used the connectivity-based parcellation procedure modified from our previous study ^33,37^. Taking an initial region as a seed mask, probabilistic tractography was applied to the native gray/white matter interface using probtrackx ^128,129^. Each vertex was sampled 10,000 times based on the orientation probability model for each voxel, with a curvature threshold of 0.2, a step length of 0.5 mm, and a number of steps of 3200. The pial surfaces were set as stop masks to prevent streamlines from crossing sulci. The connectivity of voxels that were only reached by no more than 4/10,000 streamlines was removed. This process resulted in a whole-brain connectivity profile, a matrix (*M*×*N*) between all the vertices in the ROI (*M*) and the whole-brain voxels (*N*).

A cross-correlation matrix (*M*×*M*) was calculated to quantify the similarity between the connectivity profile of vertices in the ROI. Then, the cross-correlation matrix across all subjects was averaged to construct a group matrix. Spectral clustering was used to define the distinct subregions ^37^. Since the number of clusters must be predefined, we explored 2-12 parcellations. After relabeling across subjects using a group-level labelling scheme as the reference ^130^, we used cross-validation to determine the optimal solution that yielded the greatest consistency across the subjects considering three clustering indices: Cramer’s V (CV), Dice coefficient (Dice), and topological distance (TpD). The details of the description of these indices can be found in a previous study ^131^. The local peak of Cramer’s V and Dice indicates better reproducibility than the surrounding solutions, and a TpD closer to 0 indicates a more similar topological arrangement between hemispheres. The pairwise strategy was used to evaluate the reproducibility of the parcellation. All pairs of subjects were evaluated, and the average consistency was computed. The optimal number of clusters was selected based on searching for the local peak of Cramer’s V or Dice primarily, as well as smaller TpD.

The subcortical regions were parcellated similarly but were conducted in volumetric space. For a broader application, the parcellation of the cortex was also mapped to the volumetric space by assigning each gray matter voxel to the label of its closest vertex.

#### Comparisons with existing parcellations

Previous parcellations of chimpanzee brains were used to compare with ChimpBNA, including (1) The initial seed parcellation of ChimpBNA (Init); (2) Bailey and Bonin’s brain atlas of chimpanzees, which has a digital 3D version (BB38) ^40^; (3) Davi130 parcellation (Davi130) ^12^.

We compared the parcellations using a homogeneous metric, the distance-controlled boundary coefficient (DCBC) ^50^. The basic idea underlying DCBC is that any two vertices/voxels within the same region should have more similarity between their structural connectivity profiles than those belonging to distinct regions. Taking the spatial smoothness of brain data into consideration, DCBC binned each vertex/voxel pair (bin step = 1 mm) based on their geometric distance ranging from 0 to 100 mm and then compared the similarity between the within- and between-parcel within each bin. A higher DCBC indicates that the parcellation reflected a more homogeneous topography based on structural connectivity. The DCBC of ChimpBNA was compared with that of other parcellations across subjects using paired t-test and adjusted using Bonferroni correction ^132^.

#### Mapping anatomical connectivity patterns

To compute the structural connectome between all subregions, we performed probabilistic tractography sampling 10,000 times for each voxel of each subregion and counted the connected streamlines as the weight of the edge between the subregions of each subject. The connectome of subjects was normalized and averaged to generate a group connectome for subsequent analysis.

#### Hemispheric asymmetry analysis

The surface area and gray matter volume of each subregion in ChimpBNA were computed from FreeSurfer. The hemispheric asymmetry of subregions was determined using the formula *Asym* = (*L* - *R*) / (*L* + *R*) * 0.5 ^12^, where the *L* and *R* represent the structural index for each subregion in the left and right hemispheres, respectively. Positive *Asym* values indicate a leftward asymmetry, and negative values indicate a rightward asymmetry. One-sample t-test were conducted for each subregion, and significant leftward or rightward asymmetry was determined with a *p* < .05 after correcting for multiple comparisons using Bonferroni correction ^132^.

### Connectivity Analysis between Species

#### Construction of connectivity blueprints

Using XTRACT ^53^, we reconstructed 45 homologous white matter tracts ^44^ for each human and chimpanzee subject. We performed probabilistic tractography using FSL’s probtrackx2 ^128,129^ to map the whole-brain connectivity pattern. Specifically, the white surface was set as a seed region tracking to the rest of the brain with the ventricles removed. The pial surface was used as a stop mask to prevent streamlines from crossing sulci. Each vertex was sampled 10,000 times (10,000 trackings) based on the orientation probability model for each voxel, with a curvature threshold of 0.2, a step length of 0.5 mm, and a number of steps of 3200. This resulted in a (*whole-surface vertices*) × (*whole-brain voxels*) matrix for further analysis.

Tractography of the white matter tracts was vectorized to a (*tracts*) × (*whole-brain voxels*) matrix and was multiplied by the whole-brain connectivity matrix, deriving a (*tracts*) × (*whole-surface vertices*) matrix, i.e., connectivity blueprint ^45^. We averaged the connectivity profiles of the vertices within each subregion of the HumanBNA and ChimpBNA to form the regional connectivity blueprints across subjects. The matrix columns showed each subregion’s connectivity distribution pattern, while the rows provided the cortical termination patterns of the tracts. Note that the middle cerebellar peduncle (MCP) does not project to the cortex. We disregarded the tract when constructing blueprints. So the final *tracts* = 44.

#### Connectivity divergence between species

We used connectivity blueprints built for each species to explore the connectivity divergence across species ^45^. We calculated the symmetric Kullback Leibler (KL) divergence for each pair of subregions in ChimpBNA and HumanBNA and searched for the minimal value for each subregion in the HumanBNA. Specifically, for an example subregion in HumanBNA (Figure S34B, a), a connectivity blueprint (Figure S34B, b) represents how it was connected to each white matter tract. Using the KL divergence measure, this connectivity profile was compared with that of all chimpanzee subregions (Figure S34B, c). The subregion with the connectivity blueprint most similar to the human one, i.e., with the smallest KL divergence, was picked, and this KL divergence value was assigned to the human subregion (Figure S34B, d). A higher value on this cross-species divergence map means that the subregion has a more dissimilar connectivity profile absent in chimpanzees.

#### Correlation with expansion maps of surface areas

The expansion map of surface areas between chimpanzees and humans was obtained from a previous study ^13^. We calculated the Spearman correlation between the cortical expansion and connectivity divergence map. Spin tests (10,000 times) were used to assess the correspondence between brain maps ^133^.

We also investigate the local correlation between these two maps. The correlation was computed using a sliding window around every vertex on the sphere, with a search kernel of 40° that corresponds to a circular window with a radius of approximately 7cm ^14^. A weighted mask derived from the multiplication of two maps was applied to the correlation map for magnification. Higher values on the local correlation map indicate marked cortical expansion and connectivity differences between chimpanzees and humans.

#### Connectivity divergence in functional networks

Resting-state functional networks were obtained from the Yeo 7-network atlas ^52^. We assigned each cortical subregion in the HumanBNA to one of the 7 networks. For each subregion in the atlas, we computed the ratio of the vertices belonging to each of the 7 networks and finally decided on the label using a majority vote. Connectivity divergence among the 7 networks was examined, and the divergence between the VIS/SMN networks and higher-order networks (the other five networks) was compared using the Mann-Whitney U test.

#### NeuroSynth term-based meta-analysis

To investigate the association between functional decoding and the resultant cross-species divergence map, we projected the meta-analytical task-based activation along the divergence map. We used 590 terms related to cognitive processes that had been selected in previous publications (the full list of terms can be found in a previous study ^134^). The activation maps of the cognitive terms were downloaded from the NeuroSynth database (https://neurosynth.org/). We generated 20 binarized masks at five-percentile increments of the divergence map, which ranged from 0-5% to 95%-100%, and input them into the subsequent meta-analysis. Each function term had a mean activation z-score per bin. Terms included in the visualization of meta-analysis results were those with the highest three z-scores in each bin. A significance threshold of z > 0.5 was added as a visualization constraint.

#### Connectivity embedding

We used connectivity blueprints as homologous features in a common space to compare atlases across species. We projected the regional connectivity blueprints of humans and chimpanzees to a low-dimensional space using t-SNE. Thus, subregions with similar connectivity profiles were grouped together in the resulting space. We also assigned subregions with labels of distinct cortical landmarks and calculated the median coordinates of cortical systems to compare across species.

#### Whole-brain level connectional lateralization

We investigated the lateralization of the connectivity patterns at the whole-brain level. Specifically, we obtained two hemispheric connectivity blueprints, *CB_L_* and *CB_R_*, which only contain left- or right-hemispheric and commissural tracts. We calculated the symmetric KL divergence between each pair of (*CB_L_*, *CB_R_*) rows ^45,53^, i.e., homotopic subregions between hemispheres. We calculated the asymmetric connectivity pattern for humans and chimpanzees separately.

#### Aligning chimpanzee and human brains into a common space

Since the species were not in a common space and could not be compared directly, we used an alignment technique based on myelin data to transform the connectivity pattern of the chimpanzee to the human cortex ^14,54^. This method implemented the registration using multimodal surface matching algorithm ^55^, initialized first by aligning a set of homologous ROIs between species, followed by aligning the whole-hemisphere myelin maps using the ROIs-registration step as initialization. The method had been shown to align the myelin maps well across species in the previous study ^14^, and the resulting transformations could account for relocations of other modalities. After the alignment, we could explore the species-shared and species-specific regions with a asymmetric connectivity patterns in a common human space. Here, we first focused on the human-specific asymmetry map, which was calculated as the asymmetric connectivity pattern map of humans minus the intersection between humans and chimpanzees ^56^, i.e., *human*-(*human*\**chimp*), with high values corresponding to regions whose connectivity patterns were asymmetric in humans but not in chimpanzees. Similarly, for the chimpanzee-specific asymmetry, we calculated it as *chimp*-(*chimp*\**human*), for the species-shared asymmetry, we calculated it as *human*chimp*.

To investigate the relationship between the species-specific asymmetry map and the connectivity divergence map we generated in the previous section, we computed a local correlation map with a weighted mask to up-weight the brain areas that both showed high values. The weighted mask was generated by multiplying the species-specific asymmetry map by the connectivity divergence map.

### Gene Analysis across Species

#### AHBA data and preprocessing

Regional microarray gene expression data were obtained from 6 postmortem brains (1 female; ages 24-57; *n* = 4 left-hemiphsere only, *n* = 2 both left and right hemispheres) provided by the Allen Human Brain Atlas (AHBA, https://human.brain-map.org/) ^57^. We only used data from the left hemisphere because few samples in the AHBA were obtrained from the right hemisphere. The data were processed using the abagen toolbox (version 0.1.1; https://github.com/rmarkello/abagen) ^135^. First, the microarray probes were reannotated to remove those that did not match a valid Entrez ID^136^. Then, the probes were filtered based on their expression intensity relative to background noise, and the probes with the maximum summed adjacency when representing the corresponding gene expression were kept, yielding 15633 genes corresponding to more than one probe. Last, we resampled the output gene expression map in fsaverage5 space to fs_LR32k space for subsequent study and then compiled the regional expression levels for each gene using HumanBNA parcellation scheme (105 subregions in the left cerebral hemisphere) ^37^ to form a 105×15633 regional transcription matrix.

#### Gene enrichment analysis

We used partial least squares regression (PLSR) to characterize the genetic mechanisms underlying the divergence map. We generated 10,000 surrogate maps using BrainSMASH ^133^ to ensure that the explained variance of the first PLS component (PLS1) was more significant than chance. 10,000x bootstrapping was used to assess the error of each gene’s PLS1 weight, and *Z*-scores were calculated as the ratio of the weight of each gene to its standard error ^137^. We obtained the genes that had *Z*-scores higher than 3 and submitted the gene sets for gene enrichment analysis. ToppGene (https://toppgene.cchmc.org/) ^138^, which contains the whole list of AHBA genes as the background gene set, was used to conduct the analysis. The following term categories were assessed: GO: Molecular Function, GO: Biological Process, GO: Cellular Component, Pathway, and Disease.

#### Cell-type enrichment analysis

We further performed cell-type enrichment analysis on the filtered genes using CellGO (http://www.cellgo.world) ^139^, a deep learning-based tool for cell type-specific gene function interpretation. We used the single-cell datasets accessible from a recent study ^113^. The single-nucleus RNA sequencing (snRNA-seq) data from postmortem brain samples from the dorsolateral PFC of 4 human adults (https://www.ncbi.nlm.nih.gov/geo/query/acc.cgi?acc=GSE207334). Six major classes of cells were provided in CellGO, including excitatory neurons (ExN), inhibitory neurons (InN), oligodendrocytes (Oligo), oligodendrocyte precursor cells (OPC), astrocytes (Astro), and microglia cells (Microglia). Subtypes of neurons were also available, including six excitatory neurons (L2/3IT, L3/5IT, L5/6NP, L6CT, L6IT, L6b) and five inhibitory neurons (LAMP5, PVALB, SST, VIP, SNCG). The enrichment P-values of the cell types resulting from the submitted genes were based on the Kolmogorov-Smirnov (K-S) test in CellGO.

#### Evolution-related analysis of genes

We first explored whether the filtered genes overlapped with human-accelerated genes related to brain processes (HAR-BRAIN genes) ^13^ and tested the significance using the hypergeometric test. Then, we quantified the evolutionary rate of these genes. The evolutionary rate, i.e., *dN*/*dS*, is defined as the ratio between the nonsynonymous substitution rate and the synonymous substitution rate ^58^. Values of *dN*/*dS* < 1, = 1, and > 1 indicate negative purifying selection, neutral evolution, and positive selection, respectively. We obtained the homologous genes between chimpanzees and humans in BioMart ^140^ and calculated the *dN*/*dS* as the chimpanzee-human evolutionary rate. Macaque data was also selected for comparison with chimpanzee-human results. Welch’s t-test was used to test the statistical significance.

#### Gene expression differences using PsychENCODE

We used comparative gene expression data from the PsychENCODE database (https://evolution.psyencode.org/) ^26^. The PsychENCODE database contains 16463 gene expressions at 16 homologous brain locations (10 cortical, 5 subcortical, and 1 limbic) in three species (human, *n* = 6; chimpanzee, *n* = 5; macaque, *n* = 5). The gene expression levels were quantified by RPKM (reads per kilobase of exon model per million mapped reads) and corrected for batch effects using the R package ComBat for normalization (v3.52.0) ^141^. We further normalized the gene expression data across cortical areas to *Z* scores within each individual ^136^ and averaged across individuals to obtain a group-level gene expression for each species. Only human and chimpanzee data was used in this study.

Due to the low spatial sampling of the cortical regions, we only selected three regions of interest that showed distinct connectivity divergence for further study, i.e., the superior temporal cortex (STC), dorsolateral frontal cortex (DFC), and primary visual cortex (V1C). The difference between the gene expression of the two species for each region was assessed using paired t-test, and the significance was determined with a *p* < .001 after correcting for multiple comparisons using Bonferroni correction ^132^. For each region, the differentially expressed genes between species were computed with the R package DESeq (v1.44.0) ^142^. The potential effects by cell-type ratio between species was investigated following the method used in a previous publication ^26^. We first selected the top 100 genes enriched in each cell type of sorted-cell of human neocortex ^143^. Genes enriched in cell type *A* were selected by ranking the genes based on the difference between mean expression in *A* and the maximum mean expression in cell types except *A*. The adjusted *p*-values of cell-type enriched genes in differential gene expression analysis between species were then examined.

## Data availability

The surface and volumetric representation files of the ChimpBNA are available at Github repo (https://github.com/FANLabCASIA/ChimpBNA). The ChimpBNA parcellation is also interactively accessible and downloadable via the web viewer (https://molicaca.github.io/atlas/chimp_atlas.html). The chimpanzee data are available at the National Chimpanzee Brain Resource (http://www.chimpanzeebrain.org/). The human data are available from the Human Connectome Project (https://db.humanconnectome.org/). The Yeo 7-network atlas can be downloaded at https://github.com/ThomasYeoLab/CBIG/tree/master/stable_projects/brain_parcellation/Yeo2011_fcMRI_clustering/1000subjects_reference/Yeo_JNeurophysiol11_SplitLabels/fs_LR32k. The human gene expression data are available in the Allen Brain Atlas (https://human.brain-map.org/static/download). The single-nucleus RNA sequencing data from the human adults are available at https://www.ncbi.nlm.nih.gov/geo/query/acc.cgi?acc=GSE207334. The gene expression data for humans and chimpanzees are available at BioMart (https://www.ensembl.org/info/data/biomart/index.html) and in the PsychENCODE dataset (https://evolution.psyencode.org/). The cell-type data is available at http://www.brainrnaseq.org/. The source data underlying Figures and Supplementary Figures are provided as Source Data at Github repo (https://github.com/FANLabCASIA/ChimpBNA).

## Code availability

The HCP-Pipeline can be found at https://github.com/Washington-University/HCPpipelines, and the NHP-HCP-Pipeline can be found at https://github.com/Washington-University/NHPPipelines. The neuroimaging preprocessing software used for the other datasets is freely available including FreeSurfer v6.0 (http://surfer.nmr.mgh.harvard.edu/), and FSL v6.0.5 (https://fsl.fmrib.ox.ac.uk/fsl/fslwiki). The gene processing pipeline is available (abagen, https://github.com/rmarkello/abagen), and the gene enrichment analysis process was conducted at https://toppgene.cchmc.org/. The cell-type enrichment analysis process was conducted at http://www.cellgo.world. The brain maps were presented using Workbench v1.5.0 (https://www.humanconnectome.org/software/connectome-workbench). The tracts were visualized using ITK-SNAP 4.0.1 (http://www.itksnap.org/) and Paraview 5.11.0 (https://www.paraview.org/).

## Acknowledgments

This work was partially supported by STI2030-Major Projects (Grant No. 2021ZD0200203), the Natural Science Foundation of China (Grant Nos. 82072099, 82202253, 62250058), and the China Postdoctoral Science Foundation (2022M722915). Data were provided in part by the National Chimpanzee Brain Resource (supported by NIH NS092988, NIH HG011641, NIH AG067419, NSF EF-2021785, and NSF DRL-2219759), Human Connectome Project, WU-Minn Consortium (Principal Investigators: David Van Essen and Kamil Ugurbil; 1U54MH091657) funded by the 16 NIH Institutes and Centers that support the NIH Blueprint for Neuroscience Research; and by the McDonnell Center for Systems Neuroscience at Washington University. The authors appreciate the English language and editing assistance of Rhoda E. and Edmund F. Perozzi, PhDs.

