## supplementary figures for "Comparative Analysis of Human-Chimpanzee Divergence in Brain Connectivity and its Genetic Correlates"

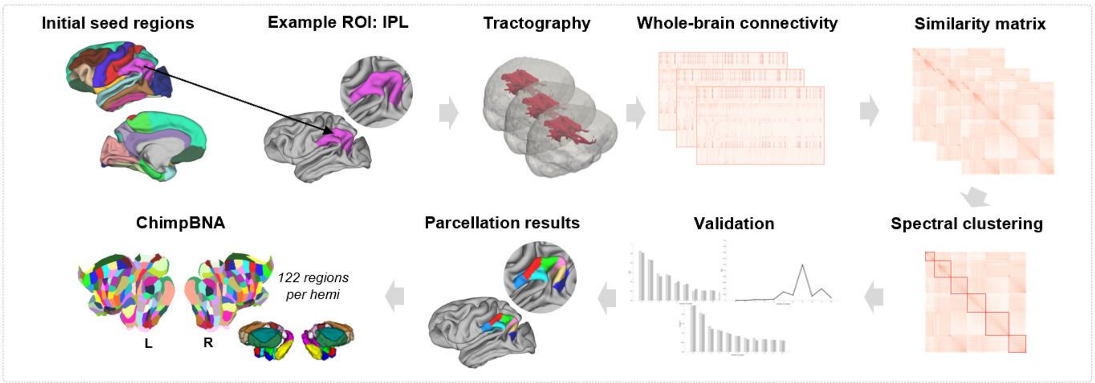


**Figure S1. Pipeline for the construction of the Chimpanzee Brainnetome Atlas (ChimpBNA).** 26 initial ROIs (19 cortical and 7 subcortical) were defined mainly based on the FreeSurfer DKT atlas. Tractography and similarity matrices were calculated for subsequent spectral clustering. The clustering results were validated using several indices, and the final parcellation of the whole brain was obtained.

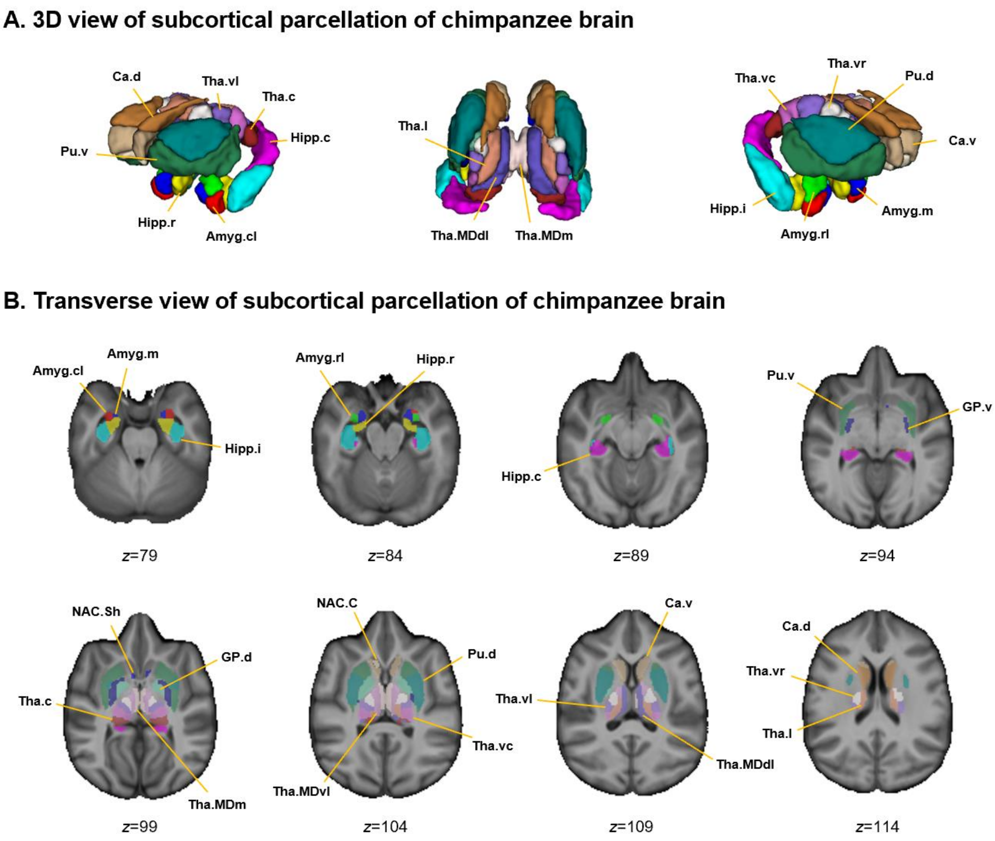


**Figure S2. Subcortical parcellation of the chimpanzee brain. (A)** 3D view of the subcortical parcellation of the chimpanzee brain. 22 subcortical subregions were identified per hemisphere using anatomical connectivity profiles. **(B)** Transverse view of the subcortical parcellation.

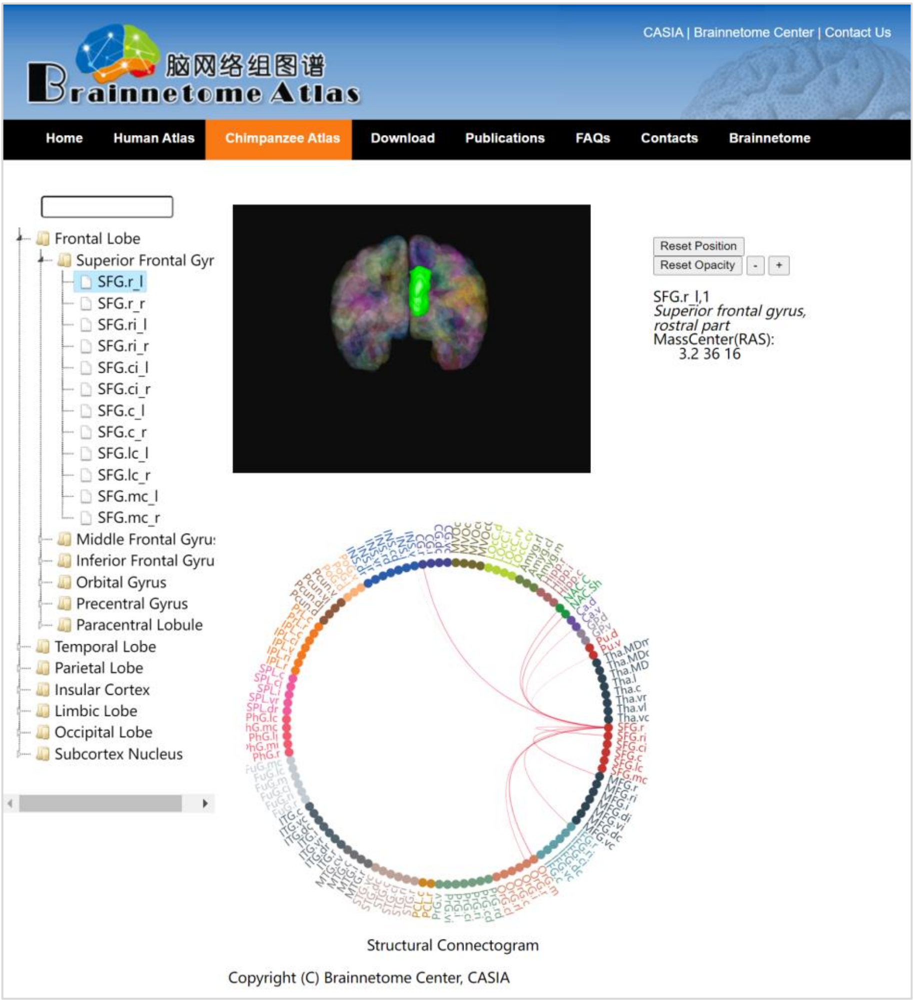


**Figure S3. Interactive website viewer of the ChimpBNA.** The website makes all the information in the ChimpBNA available to researchers. The left panel provides a hierarchical tree of the brain structures. The right panel contains a 3D atlas viewer together with a structural connectogram viewer. The website can be access at https://molicaca.github.io/atlas/chimp_atlas.html.

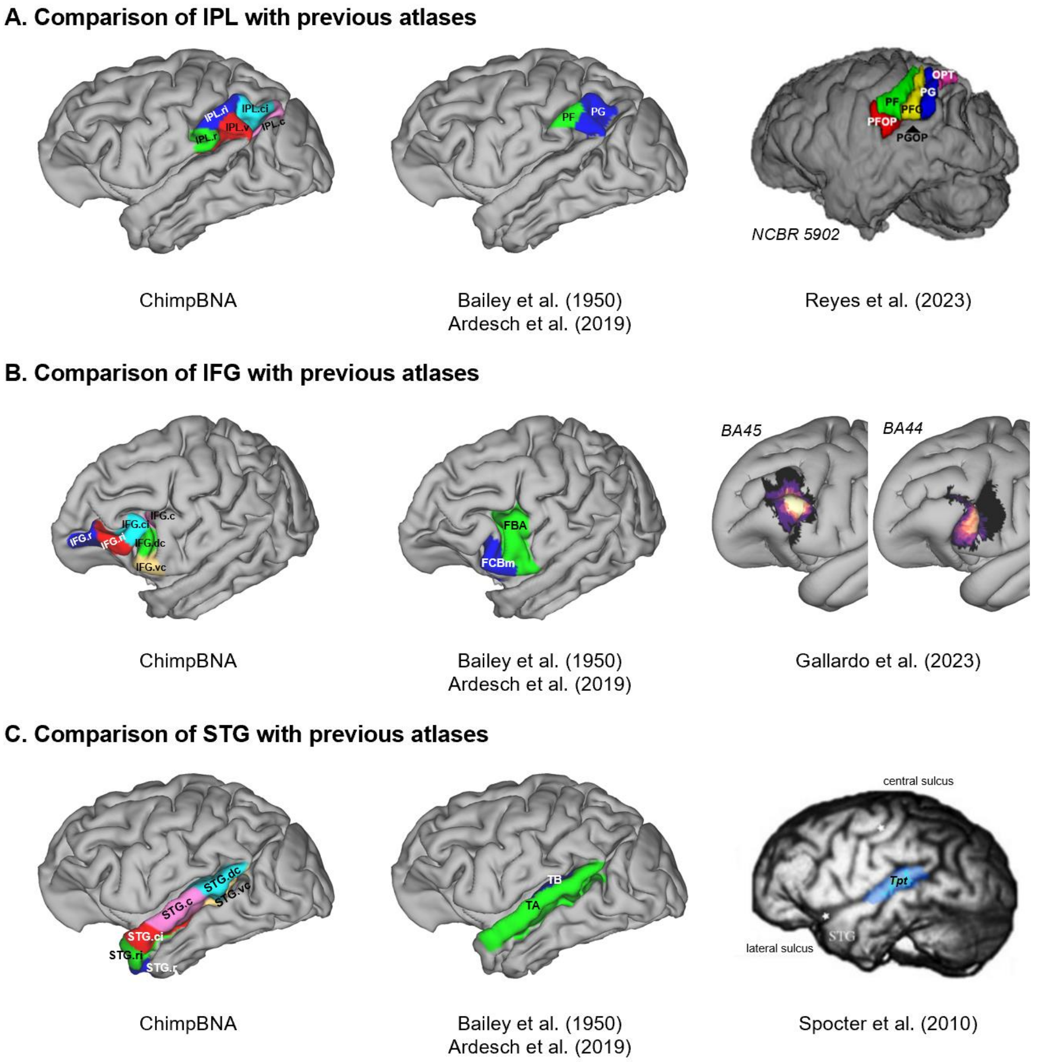


**Figure S4. Comparison of example regions with previous atlases. (A)** Comparison of IPL with previous atlases. IPL in the ChimpBNA were subdivided into 5 subregions, primarily following the rostral-caudal pattern, except for a ventral part, consistent with our previous study [1]. We note that Bailey et al. only parcellated the IPL into two subregions, i.e., a rostral one, PF, and a caudal one, PG [2]. A recent study parcellated the IPL into more refined subregions using cytoarchitecture and myeloarchitecture [3]. They identified two opercular areas and four areas on the lateral convexity of the IPL, presenting a rough agreement with the atlas we built. **(B)** Comparison of IFG with previous atlases. We parcellated the IFG into 6 subregions, two dorsal and a ventral portion of putative area 44, the rostral and caudal parts of putative area 45, and one most rostral cluster approaching the frontal pole. Since the borders of putatively homologous area 45 and 44 identified by Bailey and his colleagues, i.e., FCBm and FBA [2,4], moved back a lot, this region show incompatible parcellation.

However, Gallardo et al. confirmed the segmentation of IFG derived from published histological data [5,6]. The results showed a high consistency with our parcellation of IFG in ChimpBNA. **(C)** Comparison of STG with previous atlases. STG in the atlas by Bailey et al. was parcellated into two regions, TB and TA, which correspond to the primary auditory cortex and the remaining part of the STG [2,4]. We note that the cytoarchitectonic boundaries of area Tpt, a component of Wernicke's area which also known as BA22, was defined in a previous study [7]. The identified area Tpt was consistently located in the posterior one-third of the STG, consistent with the STG.dc in the ChimpBNA. IPL, inferior parietal lobule; IFG, inferior frontal gyrus; STG, superior temporal gyrus; STG.dc, superior temporal gyrus, dorsocaudal part. Images of all previously published parcellations were adapted from their original sources.

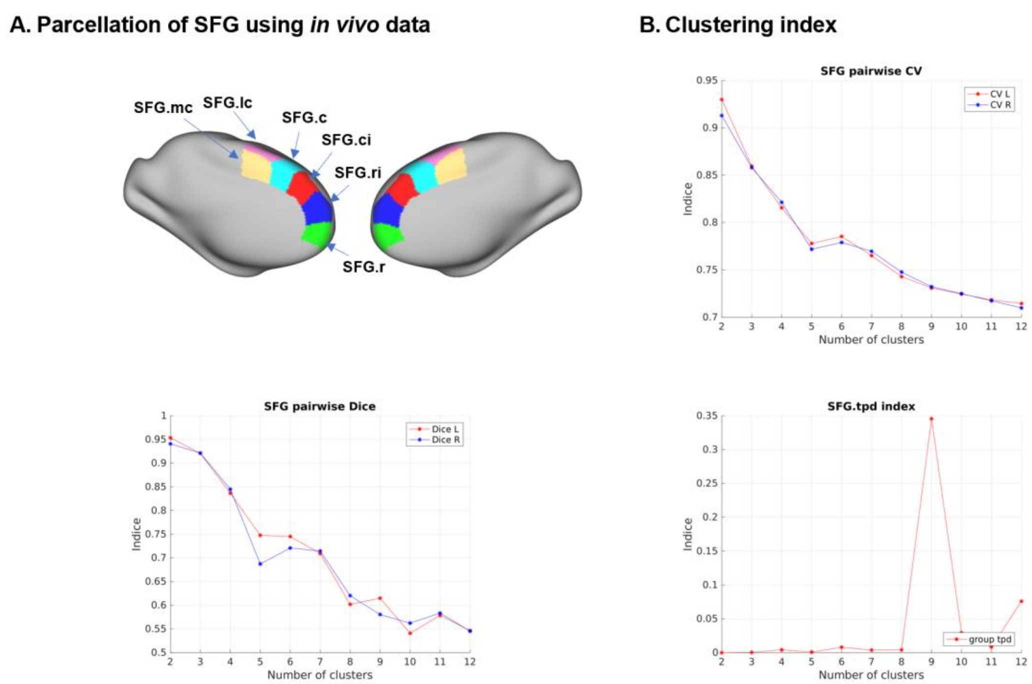


**Figure S5. Parcellation of the superior frontal gyrus (SFG). (A)** Parcellation of the SFG using *in vivo* data. **(B)** Clustering indices for the parcellation of the SFG using *in vivo* data. $k = 6$ was chosen as the final solution.

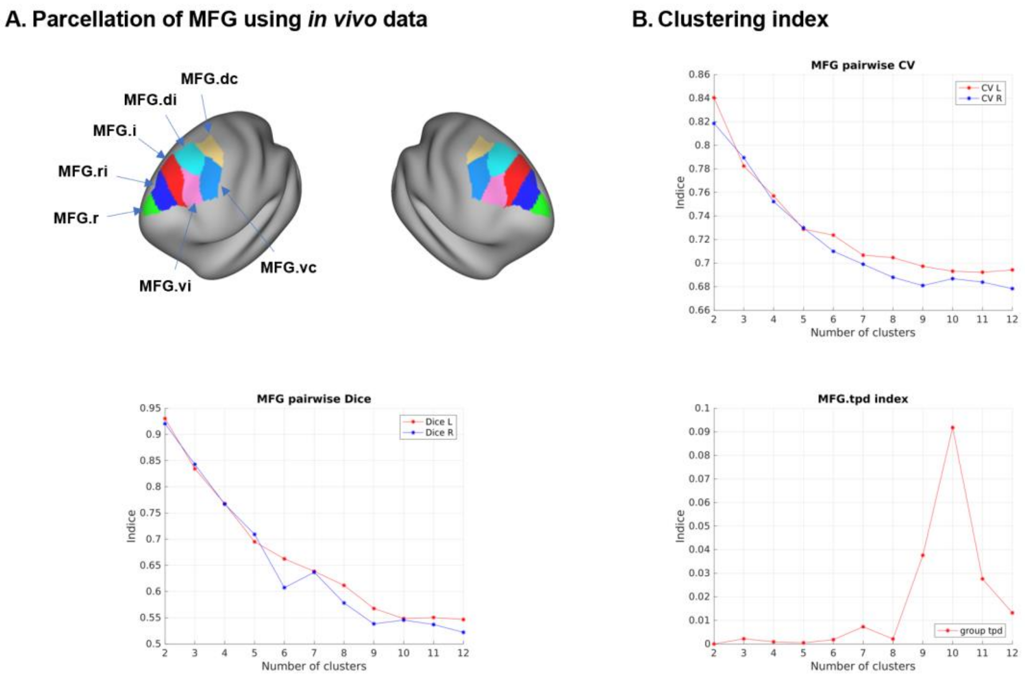


**Figure S6. Parcellation of the middle frontal gyrus (MFG). (A)** Parcellation of the MFG using *in vivo* data. **(B)** Clustering indices for the parcellation of the MFG using *in vivo* data. $k$ = 7 was chosen as the final solution.

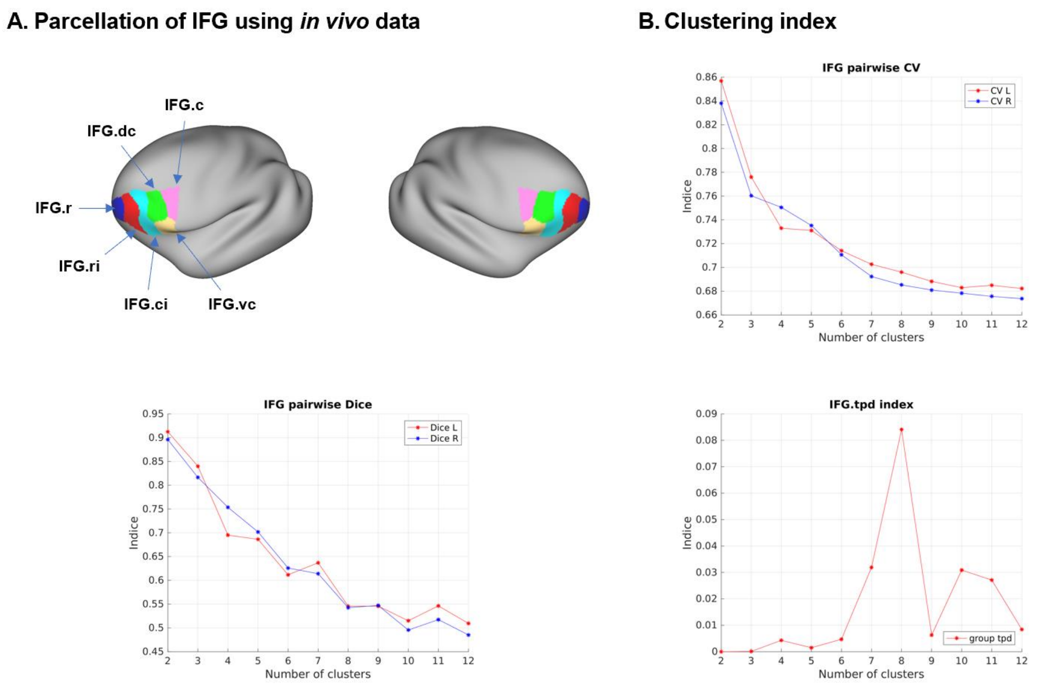


**Figure S7. Parcellation of the inferior frontal gyrus (IFG). (A)** Parcellation of the IFG using *in vivo* data. **(B)** Clustering indices for the parcellation of the IFG using *in vivo* data. $k = 6$ was chosen as the final solution.

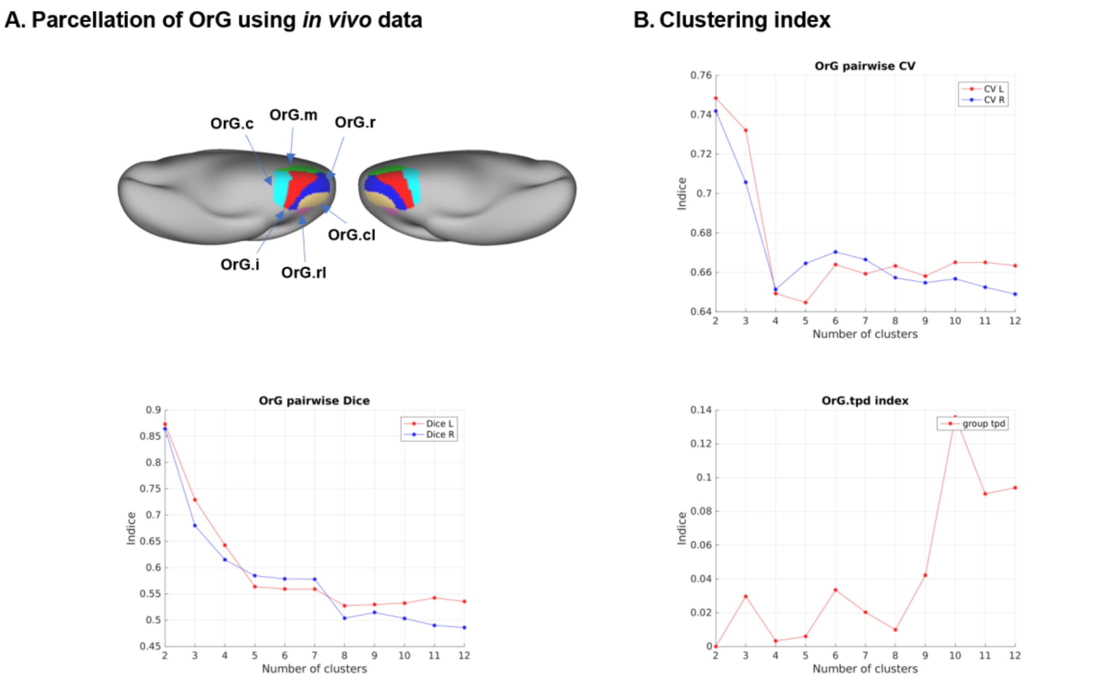


**Figure S8. Parcellation of the orbital gyrus (OrG). (A)** Parcellation of the OrG using *in vivo* data. **(B)** Clustering indices for the parcellation of the OrG using *in vivo* data. $k$ = 6 was chosen as the final solution.

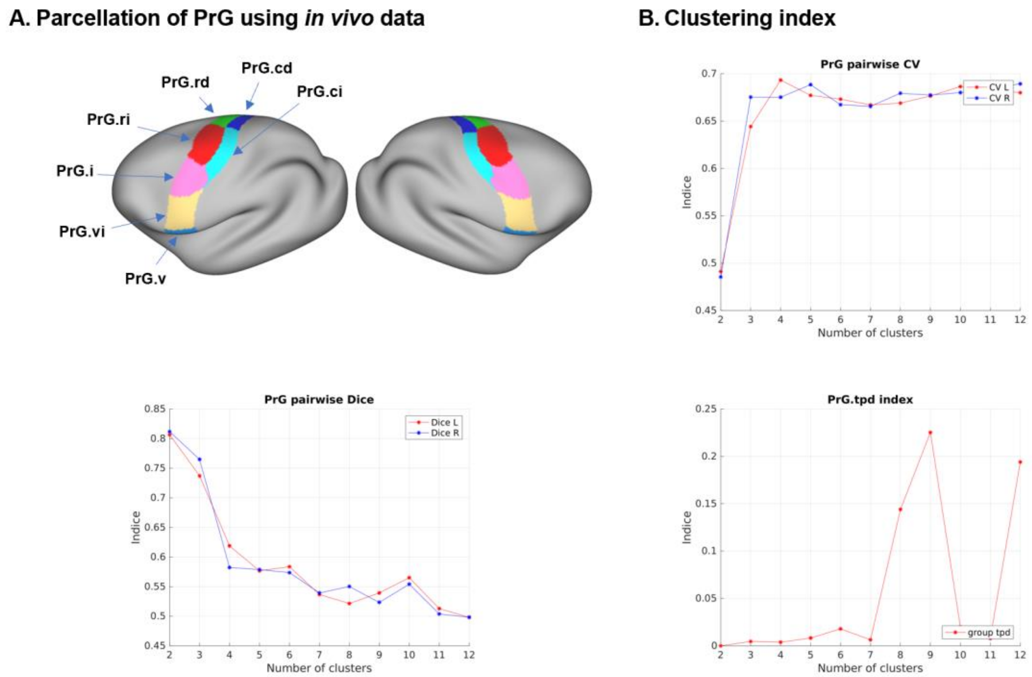


**Figure S9. Parcellation of the precentral gyrus (PrG). (A)** Parcellation of the PrG using *in vivo* data. **(B)** Clustering indices for the parcellation of the PrG using *in vivo* data. $k$ = 7 was chosen as the final solution.

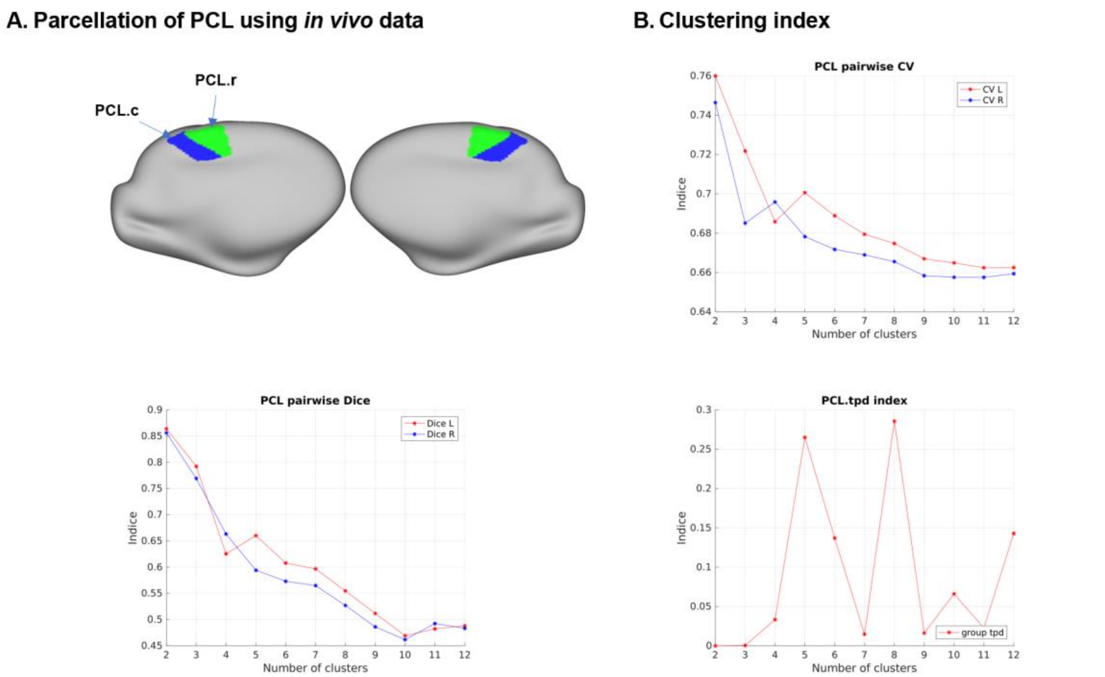


**Figure S10. Parcellation of the paracentral lobule (PCL). (A)** Parcellation of the PCL using *in vivo* data. **(B)** Clustering indices for the parcellation of the PCL using *in vivo* data. $k = 2$ was chosen as the final solution.

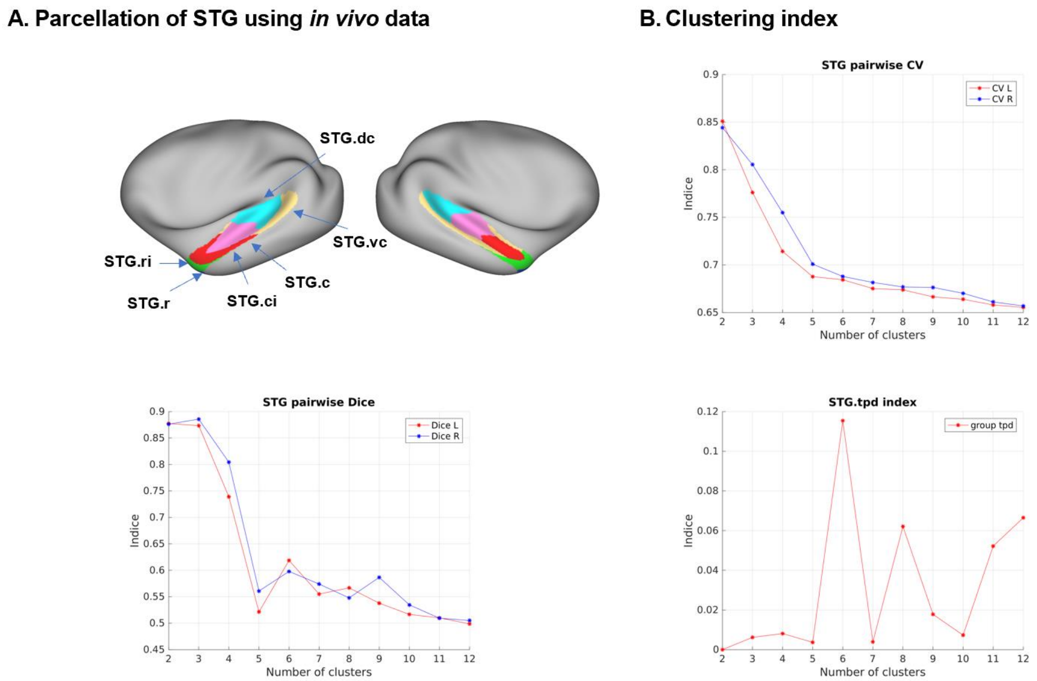


**Figure S11. Parcellation of the superior temporal gyrus (STG). (A)** Parcellation of the STG using *in vivo* data. **(B)** Clustering indices for the parcellation of the STG using *in vivo* data. $k = 6$ was chosen as the final solution.

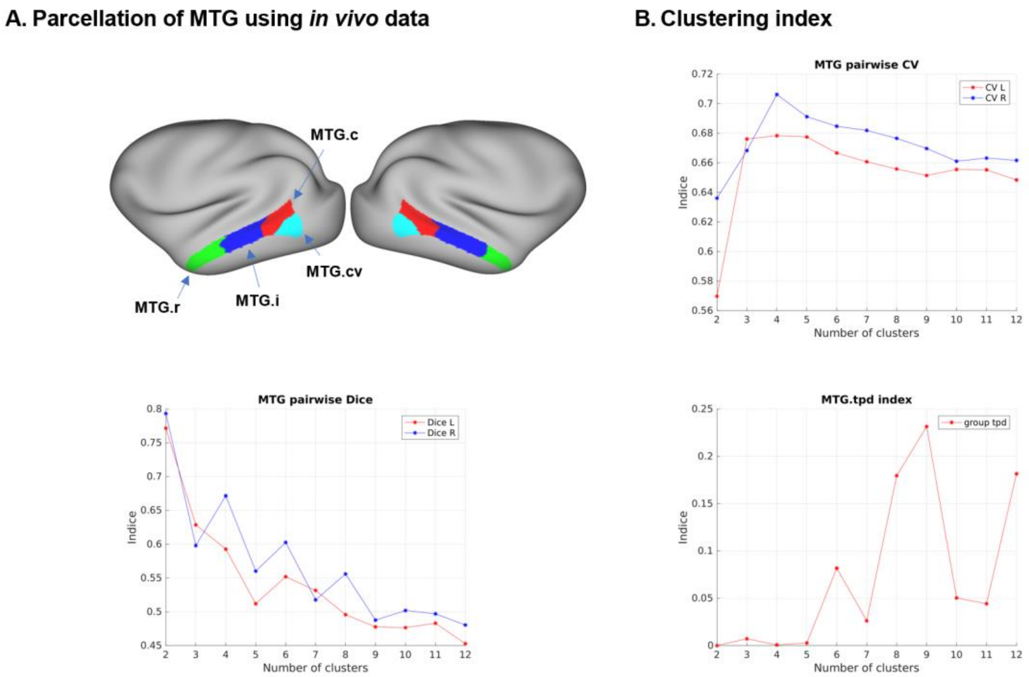


**Figure S12. Parcellation of the middle temporal gyrus (MTG). (A)** Parcellation of the MTG using *in vivo* data. **(B)** Clustering indices for the parcellation of the MTG using *in vivo* data. $k = 4$ was chosen as the final solution.

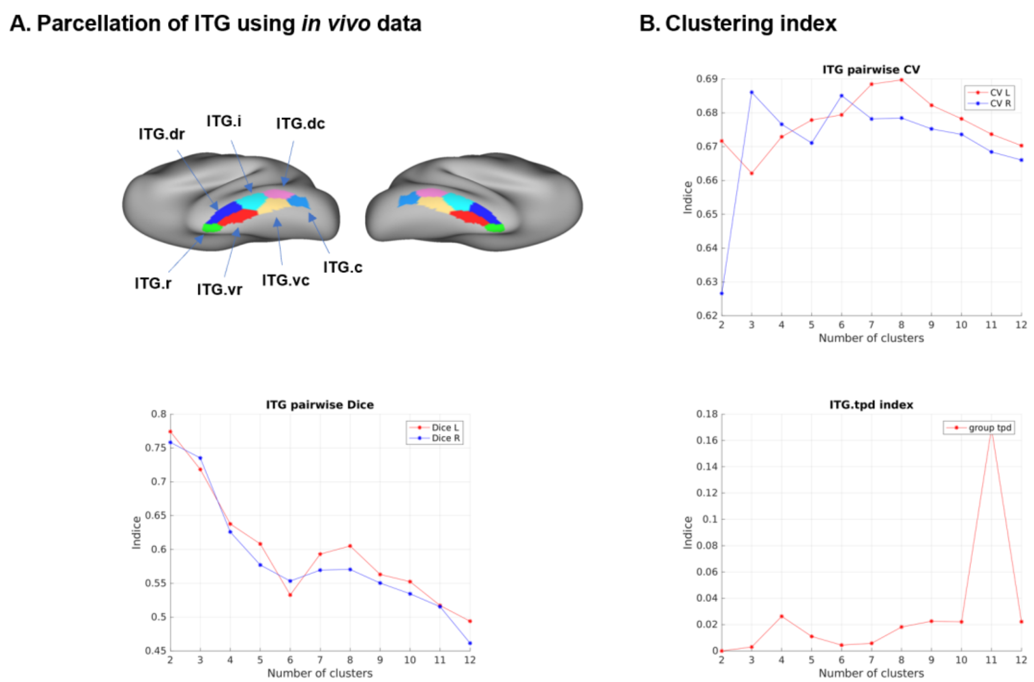


**Figure S13. Parcellation of the inferior temporal gyrus (ITG). (A)** Parcellation of the ITG using *in vivo* data. **(B)** Clustering indices for the parcellation of the ITG using *in vivo* data. $k$ = 7 was chosen as the final solution.

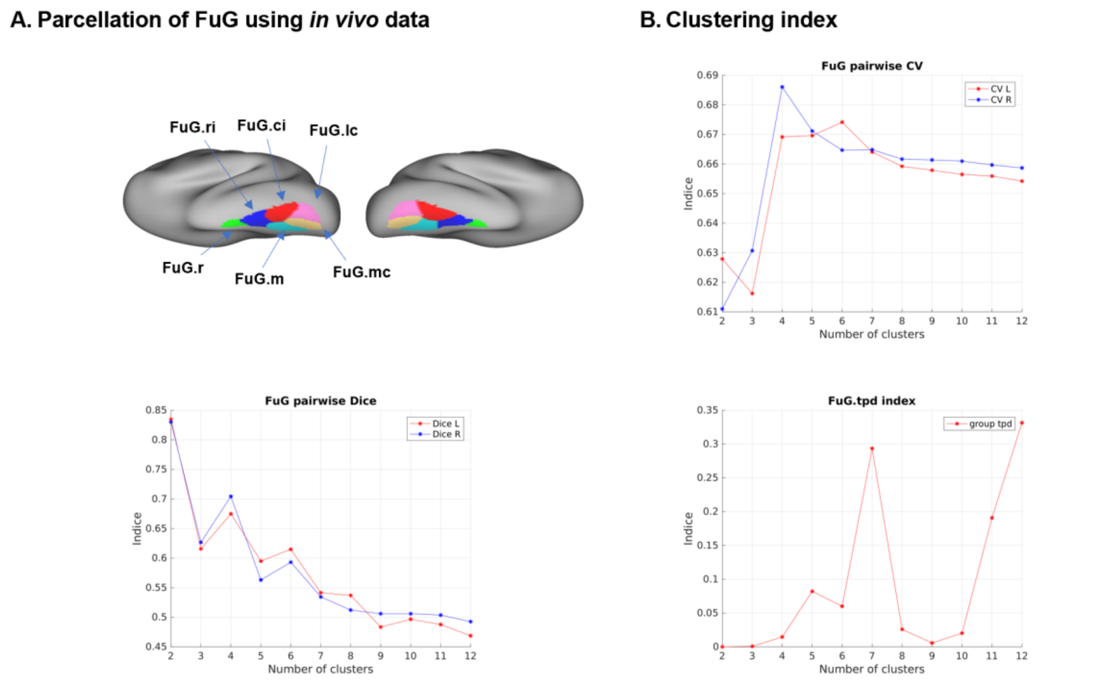


**Figure S14. Parcellation of the fusiform gyrus (FuG). (A)** Parcellation of the FuG using *in vivo* data. **(B)** Clustering indices for the parcellation of the FuG using *in vivo* data. $k = 6$ was chosen as the final solution.

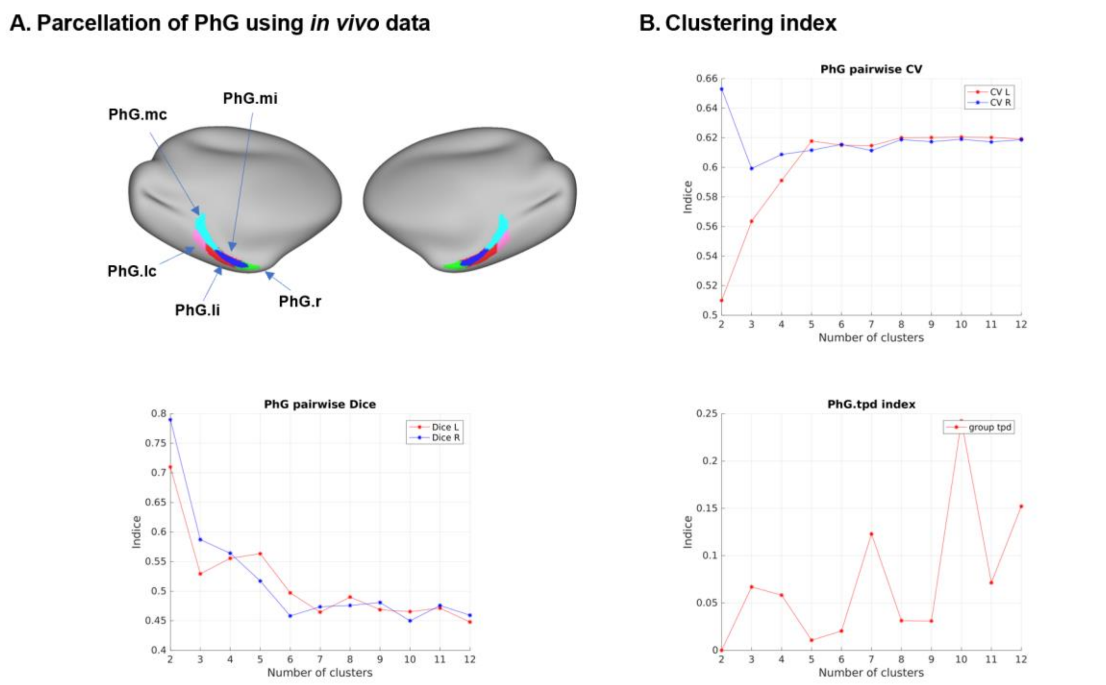


**Figure S15. Parcellation of the parahippocampal gyrus (PhG). (A)** Parcellation of the PhG using *in vivo* data. **(B)** Clustering indices for the parcellation of the PhG using *in vivo* data. $k$ = 5 was chosen as the final solution.

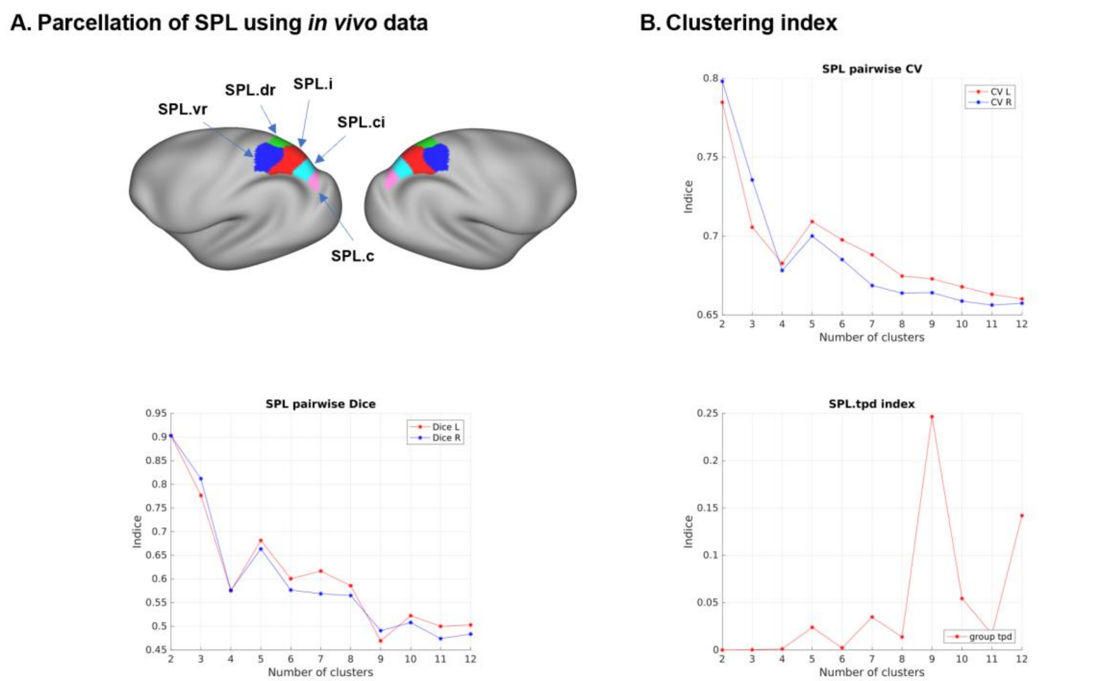


**Figure S16. Parcellation of the superior parietal lobule (SPL). (A)** Parcellation of the PCL using *in vivo* data. **(B)** Clustering indices for the parcellation of the SPL using *in vivo* data. $k$ = 5 was chosen as the final solution.

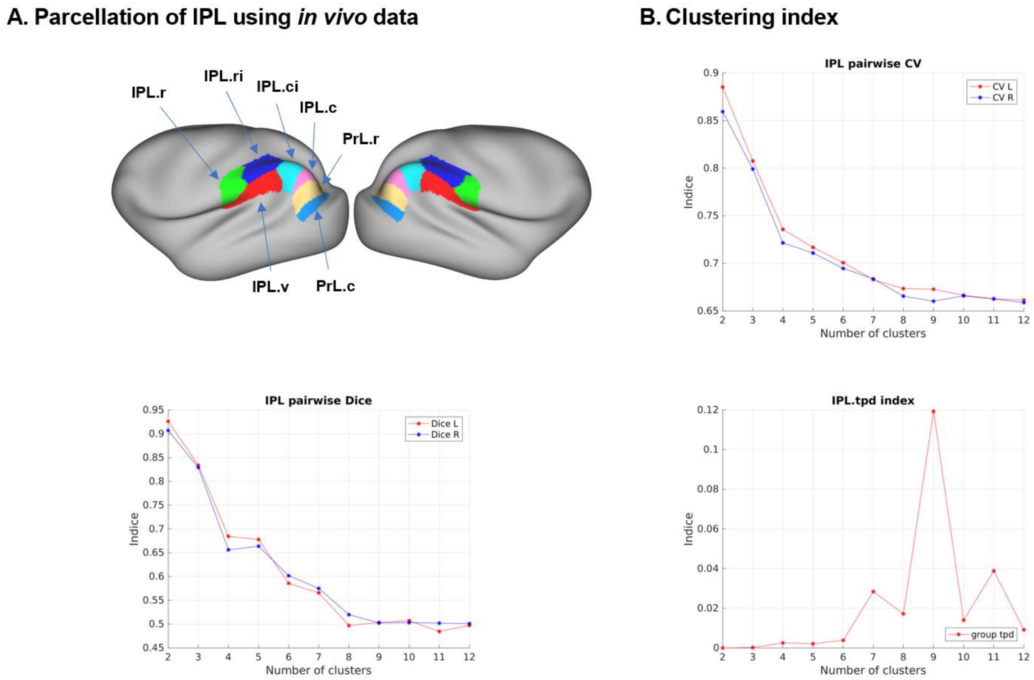


**Figure S17. Parcellation of the inferior parietal lobule (IPL). (A)** Parcellation of the IPL using *in vivo* data. **(B)** Clustering indices for the parcellation of the IPL using *in vivo* data. $k$ = 7 was chosen as the final solution. Note that the caudal parts were known as the prelunate gyrus, we additionally modified these two subregions as PrL.r and PrL.c. PrL.r, prelunate gyrus, rostral part; PrL.c, prelunate gyrus, caudal part.

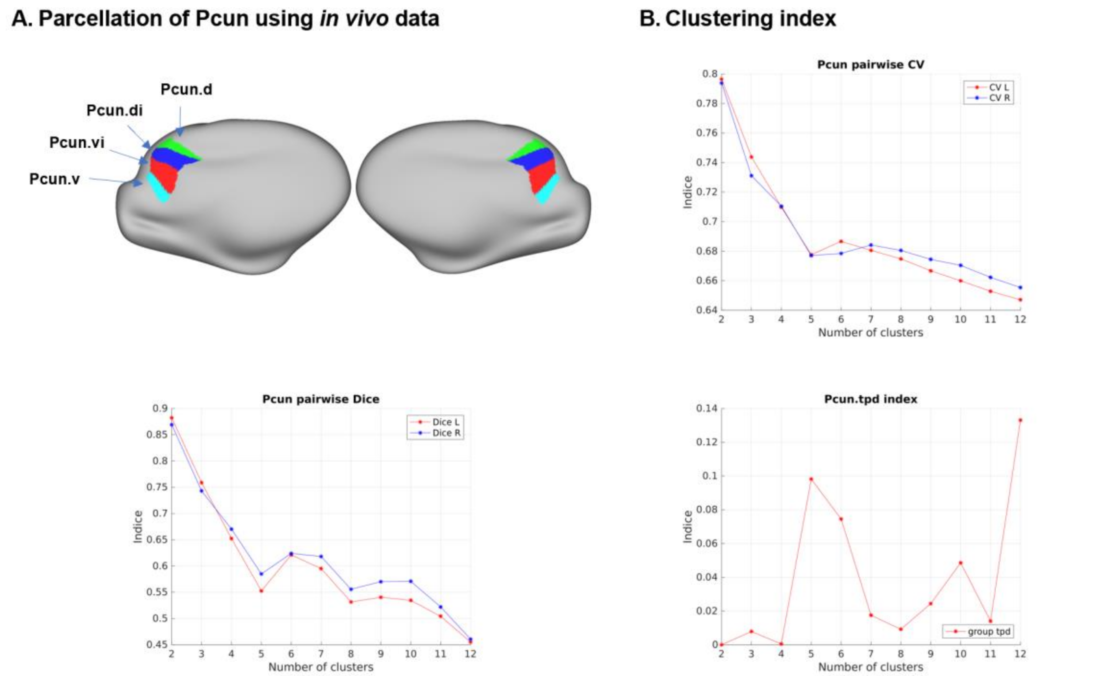


**Figure S18. Parcellation of the precuneus (Pcun). (A)** Parcellation of the Pcun using *in vivo* data. **(B)** Clustering indices for the parcellation of the Pcun using *in vivo* data. $k = 4$ was chosen as the final solution.

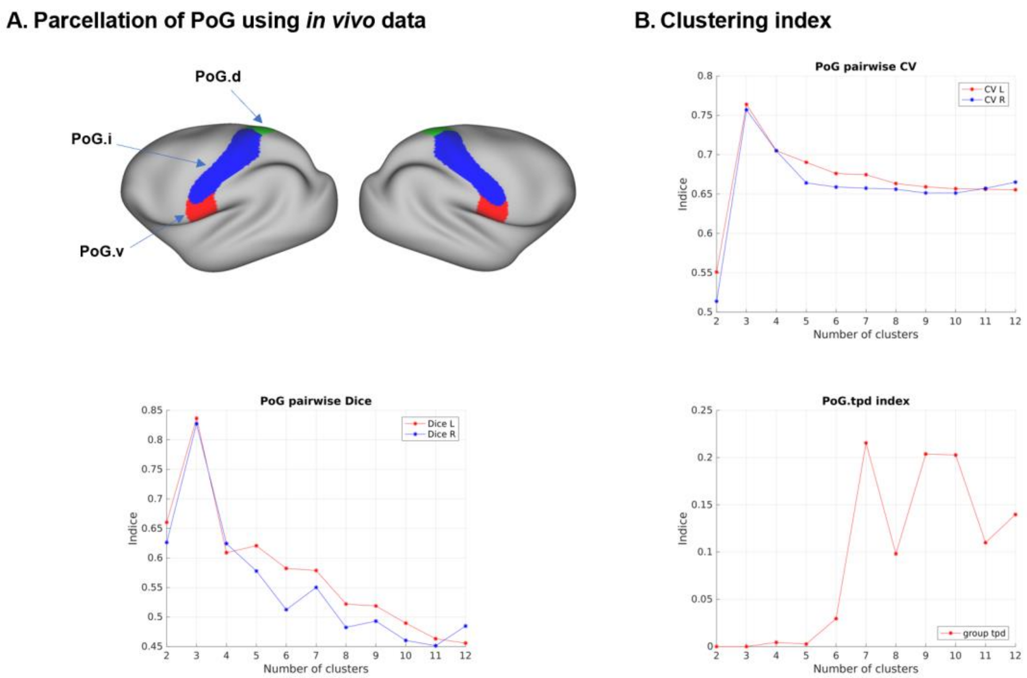


**Figure S19. Parcellation of the postcentral gyrus (PoG). (A)** Parcellation of the PoG using *in vivo* data. **(B)** Clustering indices for the parcellation of the PoG using *in vivo* data. $k = 3$ was chosen as the final solution.

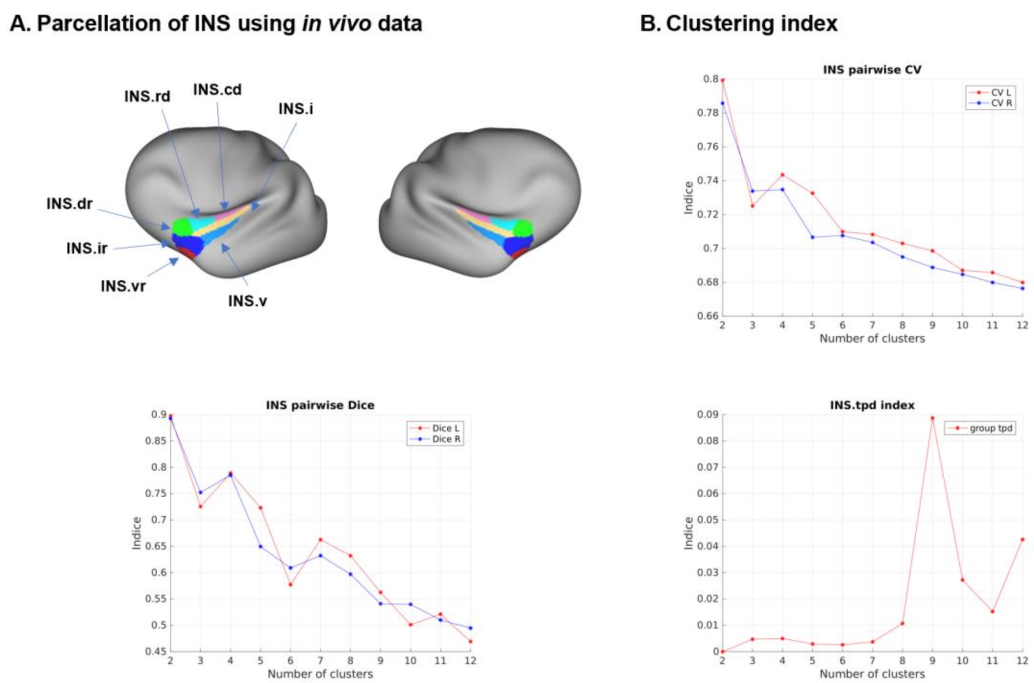


**Figure S20. Parcellation of the insular gyrus (INS). (A)** Parcellation of the INS using *in vivo* data. **(B)** Clustering indices for the parcellation of the INS using *in vivo* data. $k$ = 7 was chosen as the final solution.

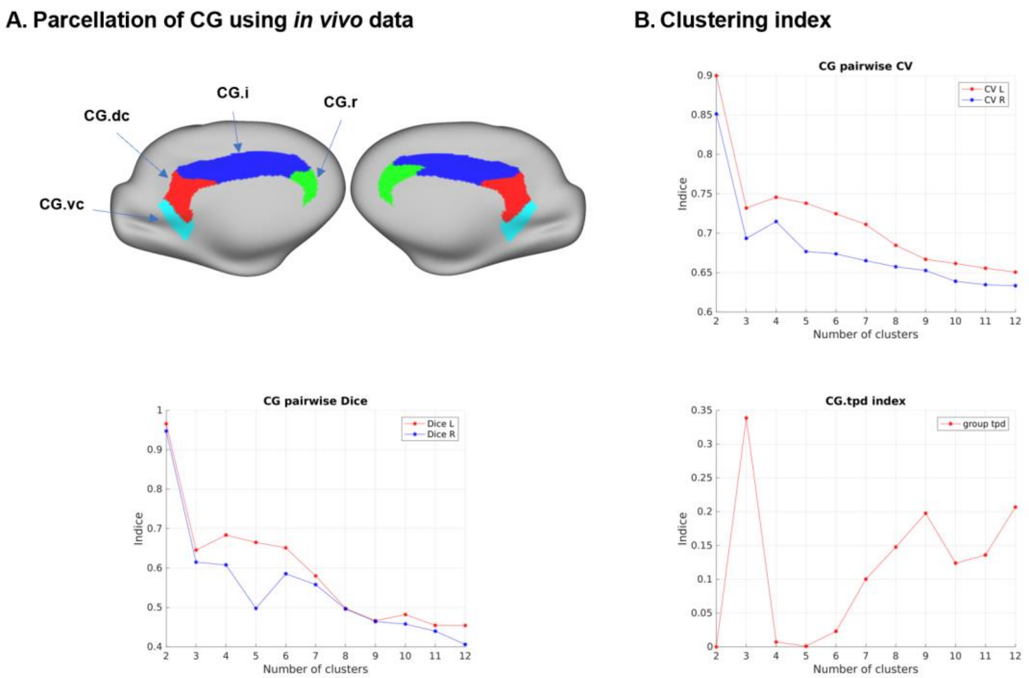


**Figure S21. Parcellation of the cingulate gyrus (CG). (A)** Parcellation of the CG using *in vivo* data. **(B)** Clustering indices for the parcellation of the CG using *in vivo* data. $k = 4$ was chosen as the final solution.

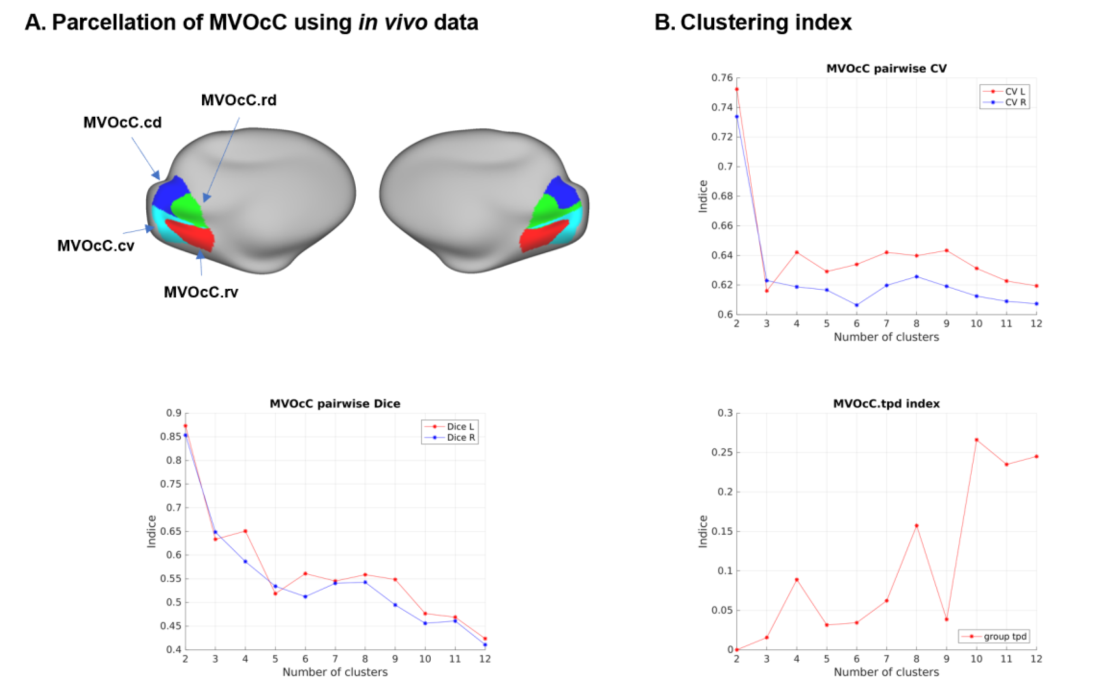


**Figure S22. Parcellation of the medioventral occipital cortex (MVOcC). (A)** Parcellation of the MVOcC using *in vivo* data. **(B)** Clustering indices for the parcellation of the MVOcC using *in vivo* data. $k = 4$ was chosen as the final solution.

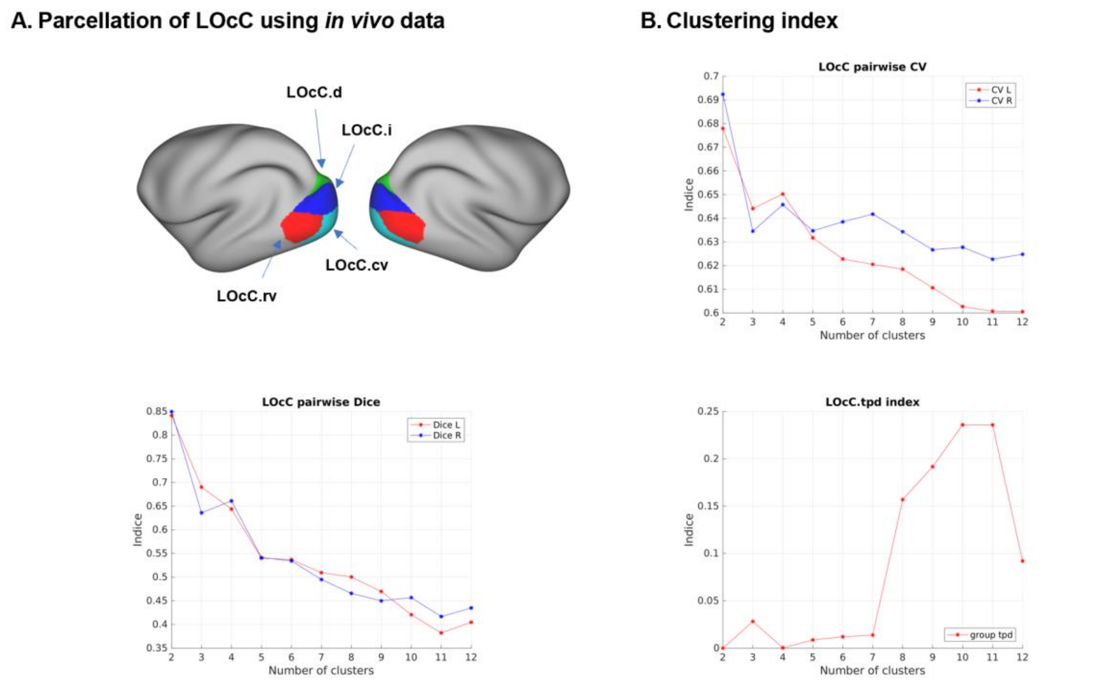


**Figure S23. Parcellation of the lateral occipital cortex (LOcC). (A)** Parcellation of the LOcC using *in vivo* data. **(B)** Clustering indices for the parcellation of the LOcC using *in vivo* data. $k = 4$ was chosen as the final solution.

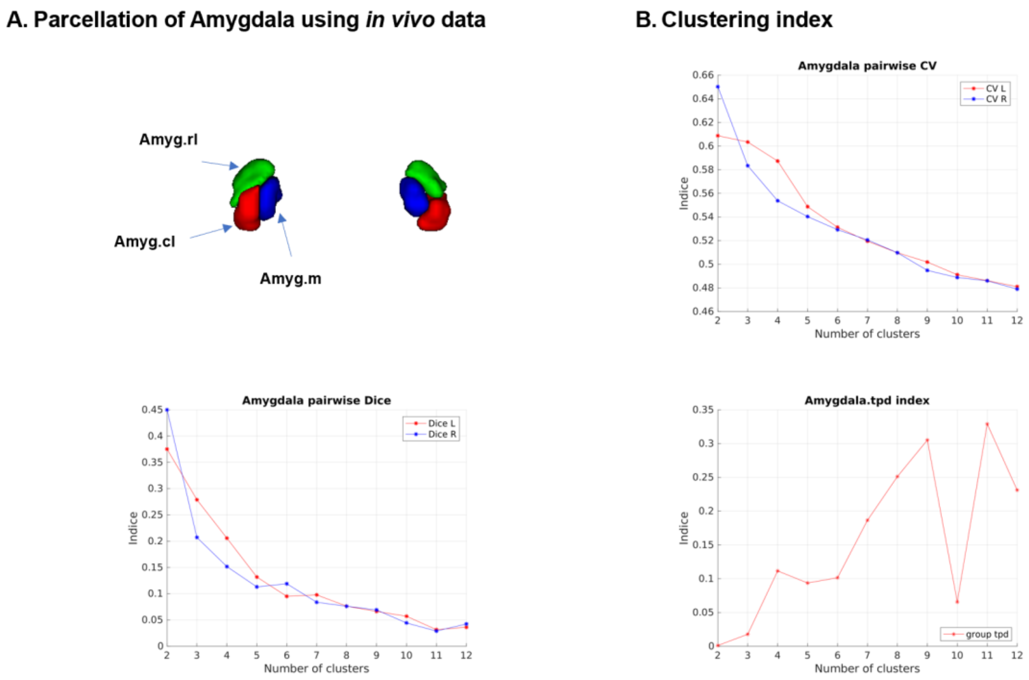


**Figure S24. Parcellation of the amygdala (Amyg). (A)** Parcellation of the amygdala using *in vivo* data. **(B)** Clustering indices for the parcellation of the amygdala using *in vivo* data. $k = 3$ was chosen as the final solution.

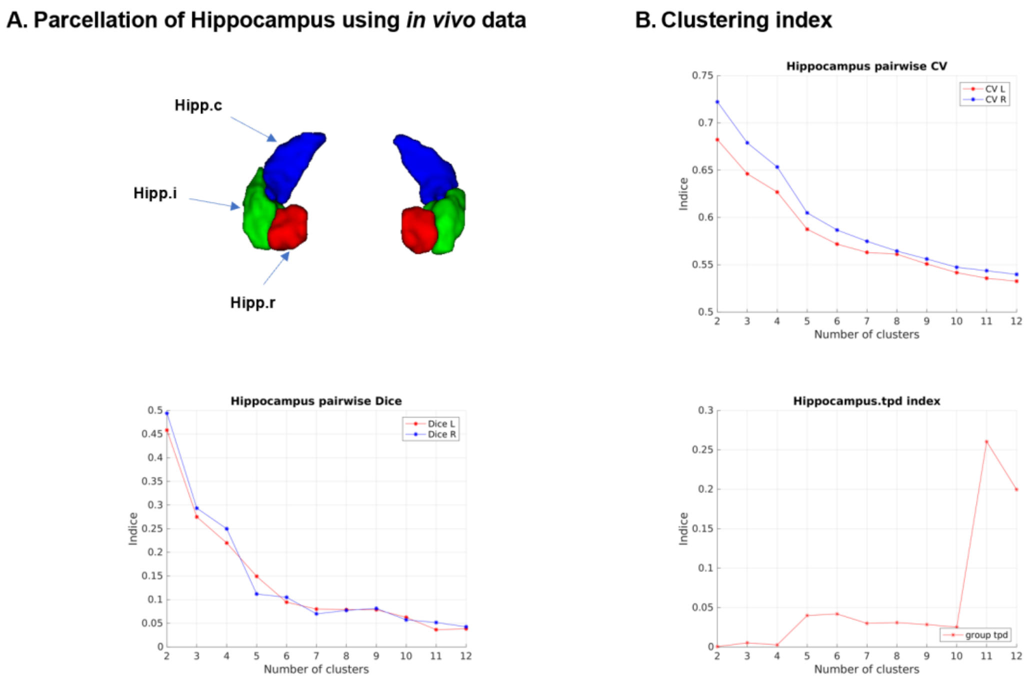


**Figure S25. Parcellation of the hippocampus (Hipp). (A)** Parcellation of the hippocampus using *in vivo* data. **(B)** Clustering indices for the parcellation of the hippocampus using *in vivo* data. $k = 3$ was chosen as the final solution.

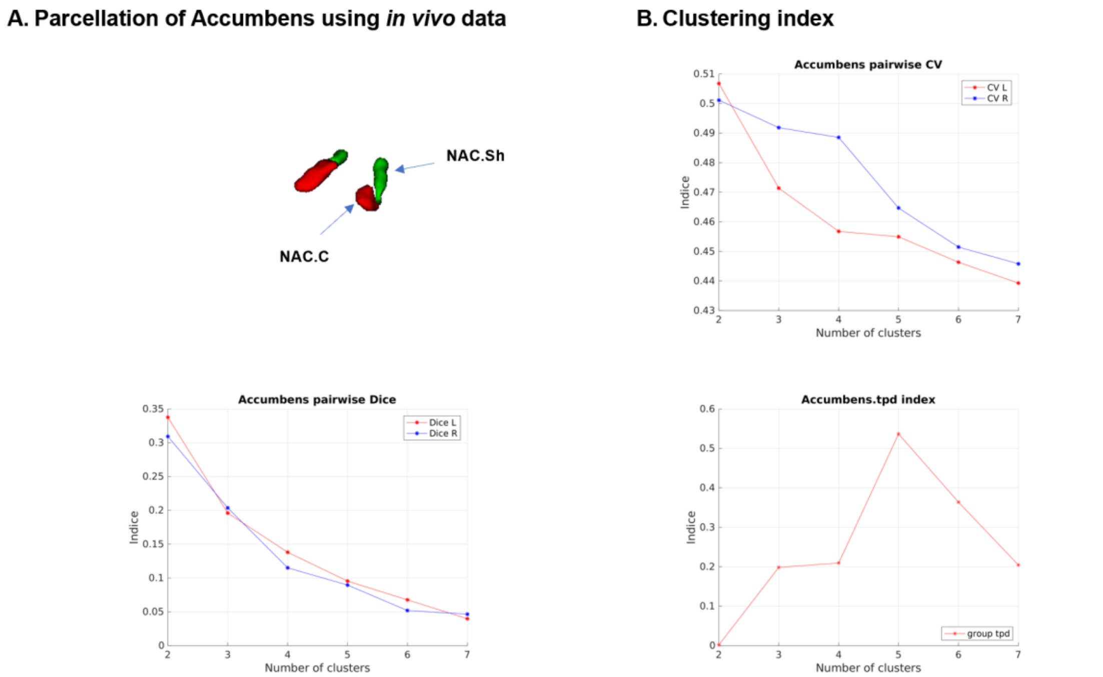


**Figure S26. Parcellation of the nucleus accumbens (NAC). (A)** Parcellation of the nucleus accumbens using *in vivo* data. **(B)** Clustering indices for the parcellation of the nucleus accumbens using *in vivo* data. $k = 2$ was chosen as the final solution.

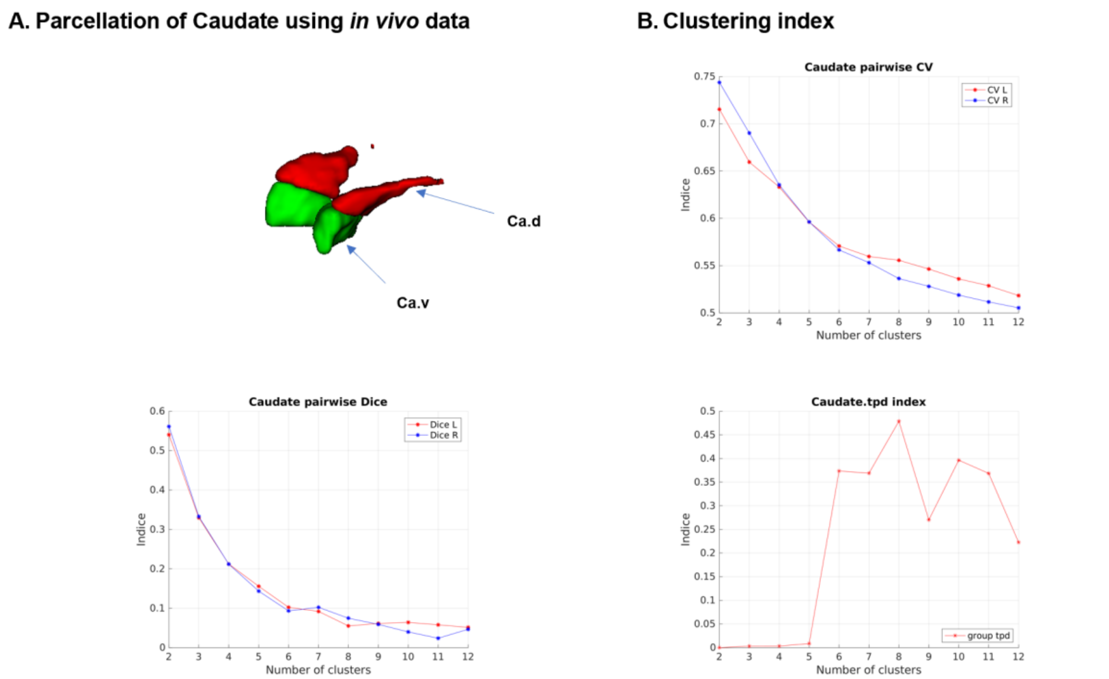


**Figure S27. Parcellation of the caudate (Ca). (A)** Parcellation of the caudate using *in vivo* data. **(B)** Clustering indices for the parcellation of the caudate using *in vivo* data. $k = 2$ was chosen as the final solution.

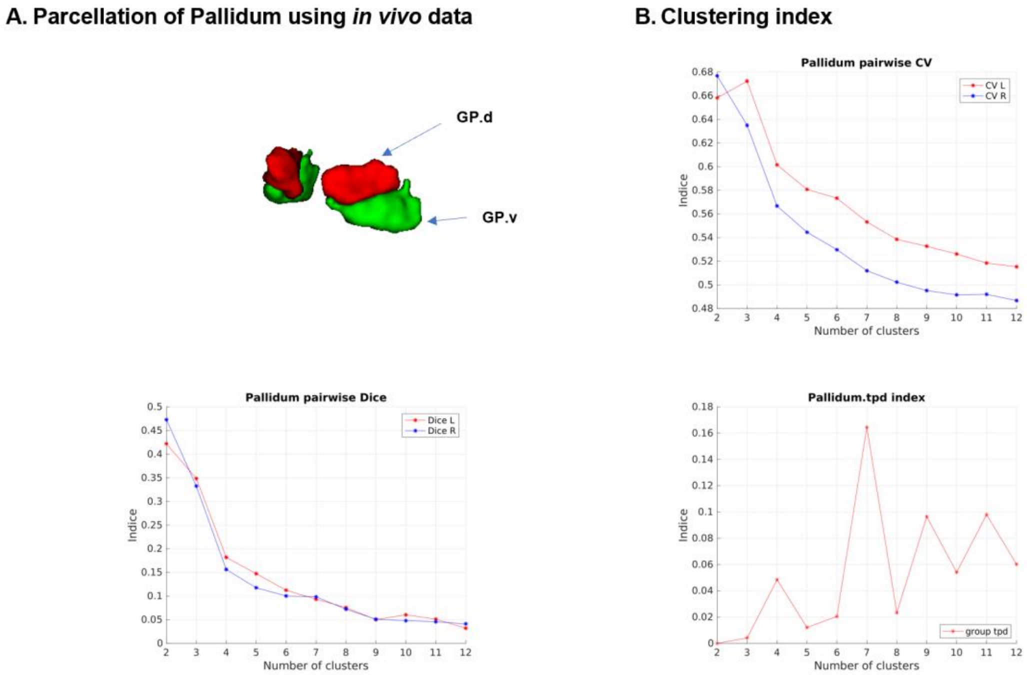


**Figure S28. Parcellation of the global pallidums (GP). (A)** Parcellation of the global pallidums using *in vivo* data. **(B)** Clustering indices for the parcellation of the global pallidums using *in vivo* data. $k = 2$ was chosen as the final solution.

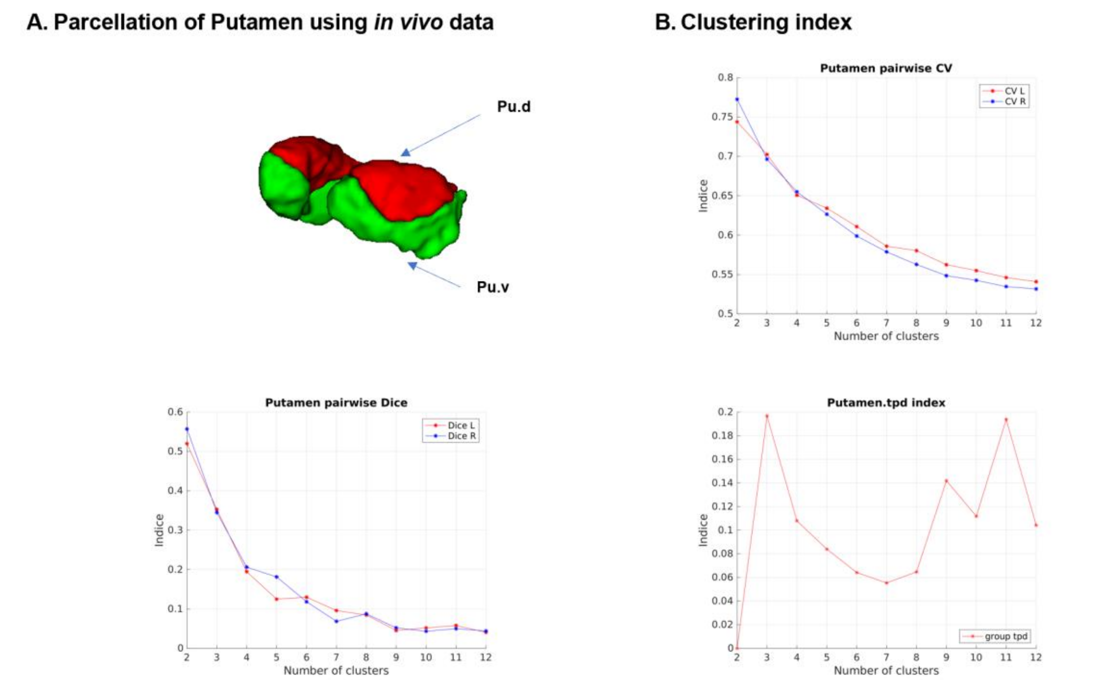


**Figure S29. Parcellation of the putamen (Pu). (A)** Parcellation of the putamen using *in vivo* data. **(B)** Clustering indices for the parcellation of the putamen using *in vivo* data. $k = 2$ was chosen as the final solution.

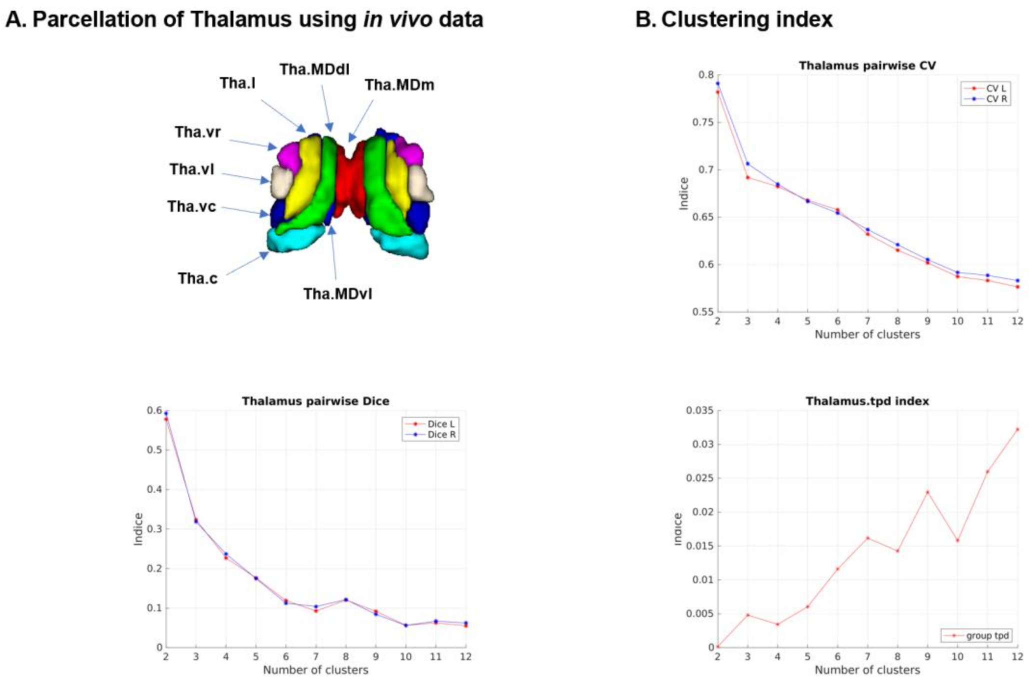


**Figure S30. Parcellation of the thalamus (Tha). (A)** Parcellation of the thalamus using *in vivo* data. **(B)** Clustering indices for the parcellation of the thalamus using *in vivo* data. $k$ = 8 was chosen as the final solution.

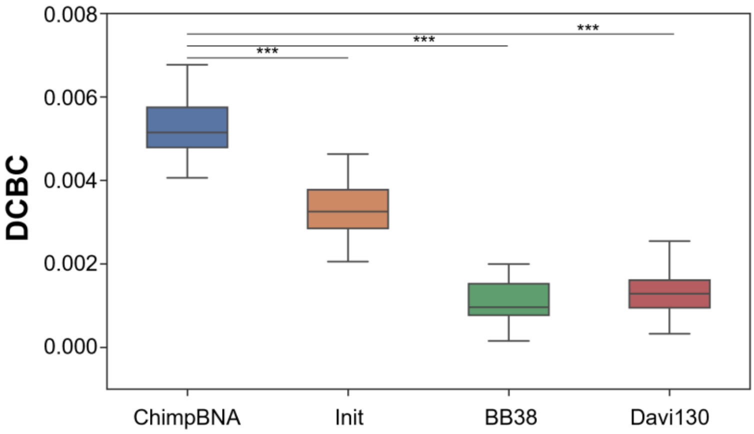


**Figure S31. Homogeneity of the ChimpBNA and other chimpanzee brain parcellations using the distance-controlled boundary coefficient (DCBC) based on structural connectivity.** The ChimpBNA showed the best performance compared with other parcellations ($p < .001$, Bonferroni corrected). *** indicates a $p$-value < 0.001.

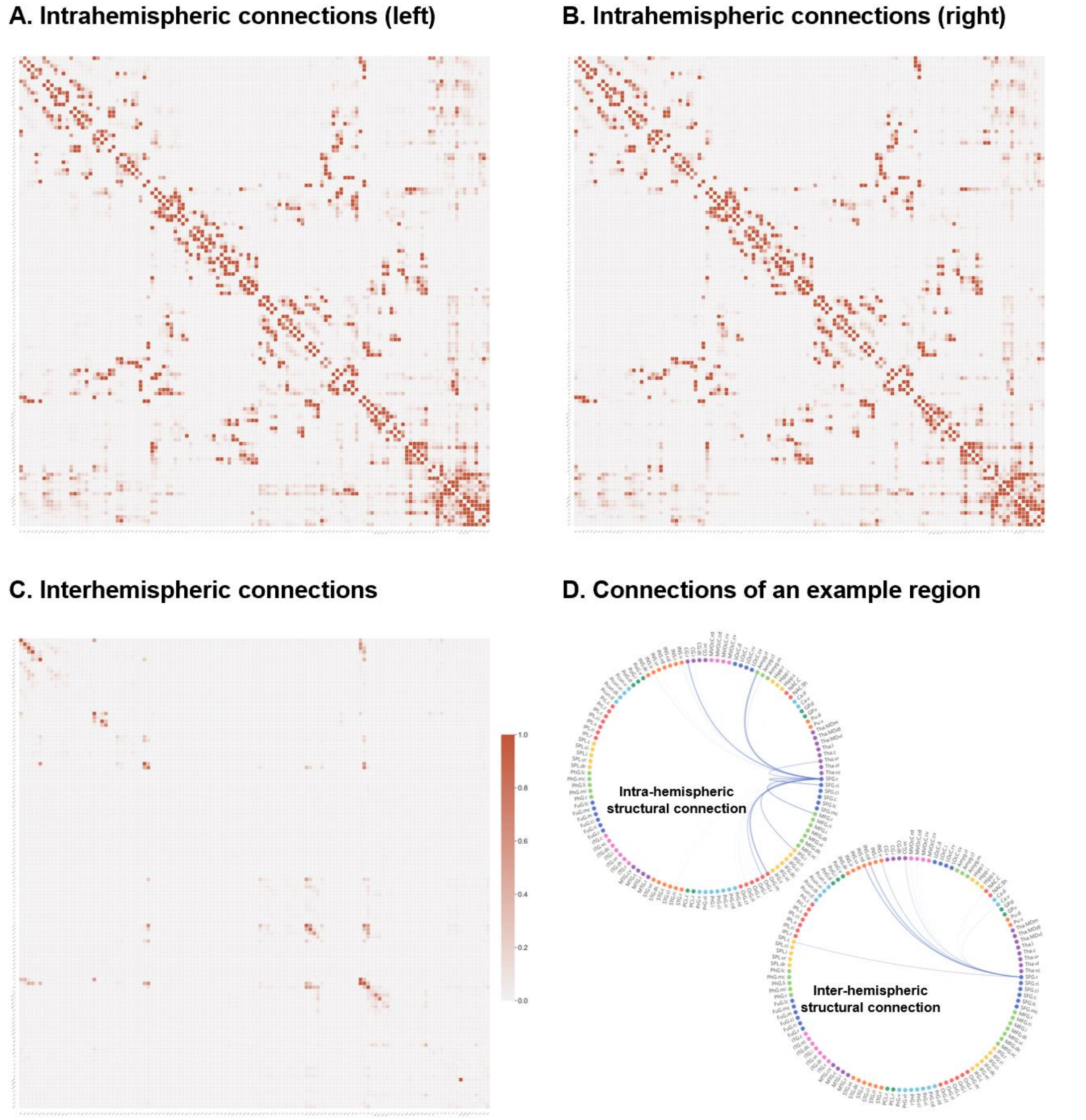


**Figure S32. Region-to-region connections of the ChimpBNA. (A)** The intrahemispheric connection matrix for the left hemisphere. **(B)** The intrahemispheric connection matrix for the right hemisphere. **(C)** The interhemispheric connection matrix across the two hemispheres. **(D)** Example of the subregional connections of the SFG.r. SFG.r, superior frontal gyrus, rostral part.

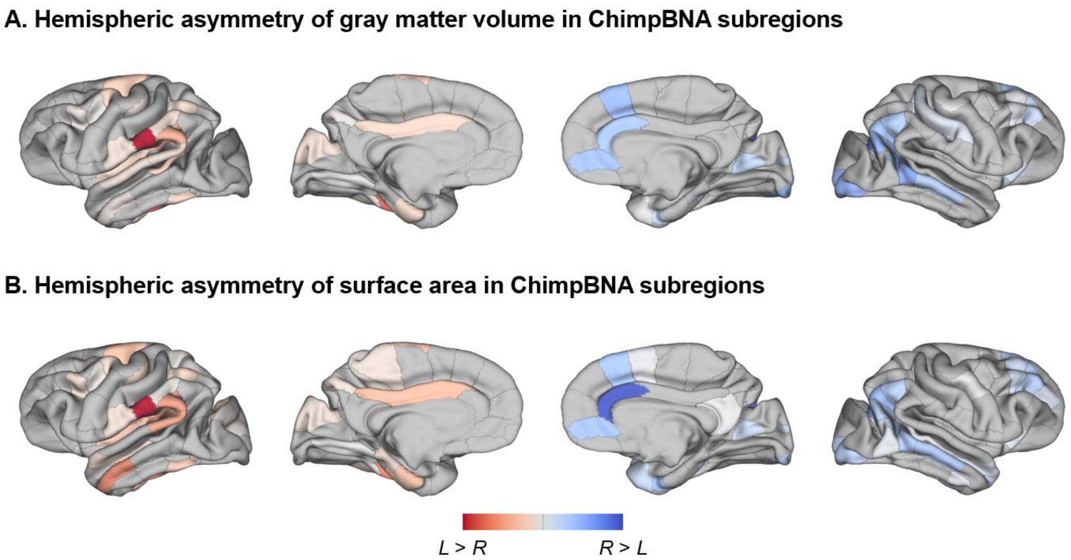


**Figure S33. Hemispheric asymmetry in Chimpanzee Brainnetome Atlas subregions. (A)** Hemispheric asymmetry of gray matter volume in ChimpBNA subregions. **(B)** Hemispheric asymmetry of surface area in ChimpBNA subregions. Red color indicates a leftward asymmetry, and blue color indicates a rightward asymmetry. The results underwent one-sample t-tests and adjusted for Bonferroni correction, $p < .05$.

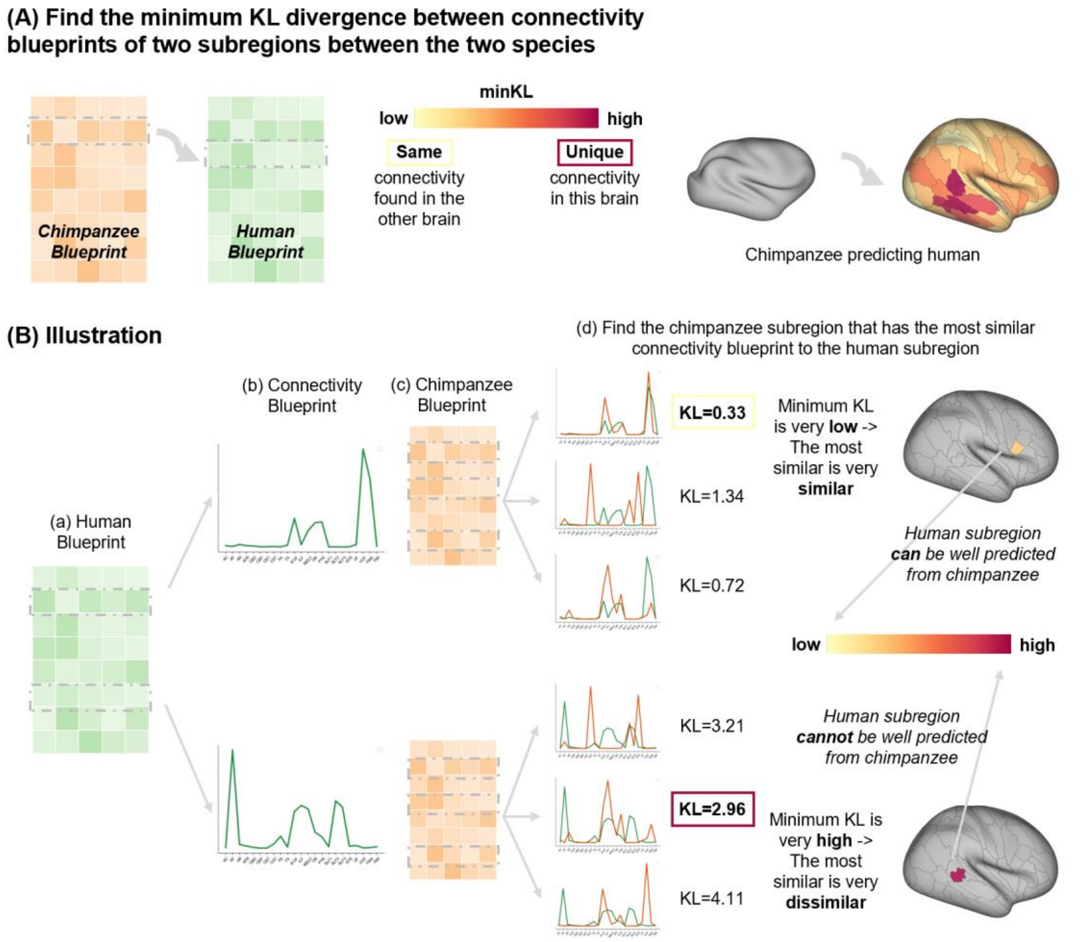


195

196 **Figure S34. Connectivity blueprint approach to investigate connectivity divergence between humans and chimpanzees. (A)** Connectional changes within the brains can be located by quantifying the dissimilarity in connectivity profiles among various regions of the two brains. **(B)** The illustration of the KL divergence computation elucidates the comparative analysis between two distinct species. For instance, within the subregion in HumanBNA (a), a connectivity blueprint (b) represents the specific connections to each white matter tract. Using the KL divergence measure, this connectivity profile was compared with that of all chimpanzee subregions (c). The subregion exhibiting the highest resemblance to the human connectivity pattern, characterized by the lowest KL divergence, was selected, with the corresponding KL divergence value attributed to the human subregion (d).

206

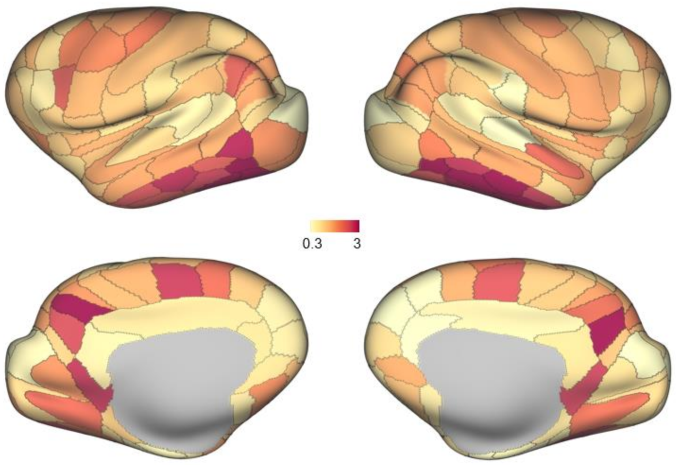


**Figure S35. Connectivity divergence between species.** Connectivity blueprints were used to calculate the KL divergence between species to determine the extent of dissimilarity between their connectivity profiles. Higher values in the divergence map indicated that the region in chimpanzees had a connectivity pattern that was more dissimilar to the regions in humans.

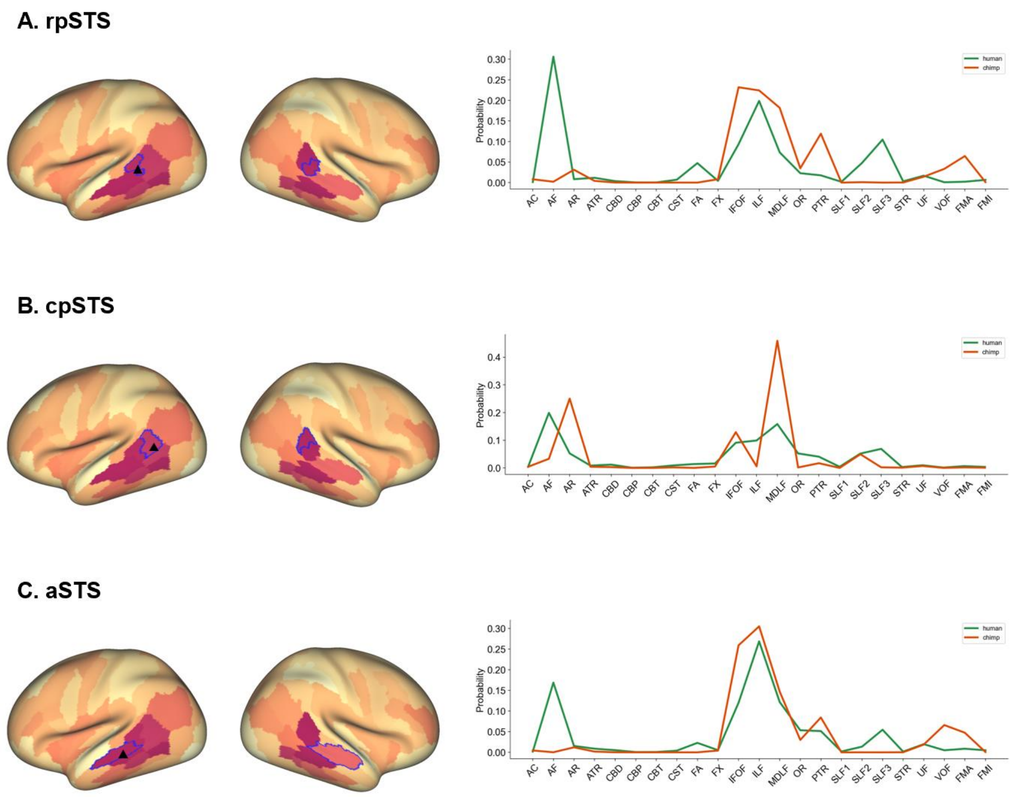


**Figure S36. Tract connectivity distribution of example subregions which showed less significant cortical expansion but higher connectivity divergence. (A)** Tract connectivity distribution of the rpSTS. **(B)** Tract connectivity distribution of the cpSTS. **(C)** Tract connectivity distribution of the aSTS. rpSTS, rostroposterior superior temporal sulcus; cpSTS, caudoposterior superior temporal sulcus; aSTS, anterior superior temporal sulcus.

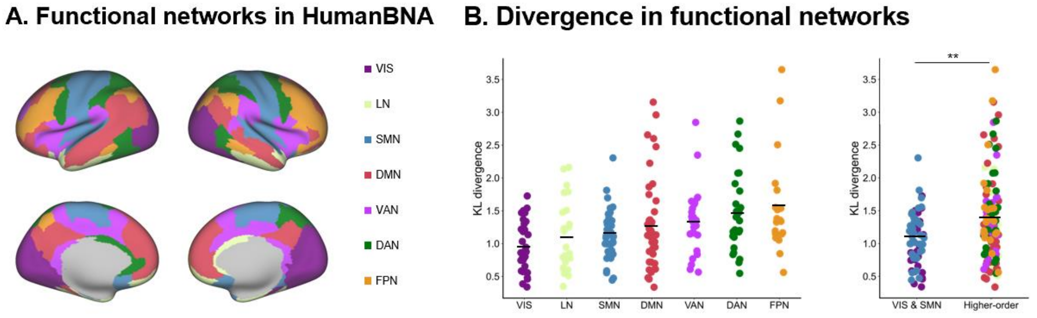


**Figure S37. Connectivity divergence in functional networks. (A)** Yeo's seven resting-state functional networks according to the HumanBNA [8,9]. **(B)** Connectivity divergence in the 7 functional networks and in higher-order cognitive networks versus VIS/SMN. Higher-order cognitive networks displayed greater divergence than the VIS/SMN (Mann-Whitney U test, $p$ = .0045), with the FPN having the greatest divergence. Dots depict cortical subregions in HumanBNA. Colors represent functional networks, as in **(A)**. Central marks are the mean KL divergence value. VIS, visual network; SMN, somatomotor network; LN, limbic network; DAN, dorsal-attention network; VAN, ventral-attention network; FPN, frontoparietal network; DMN, default mode network. ** indicates a $p$-value < 0.01.

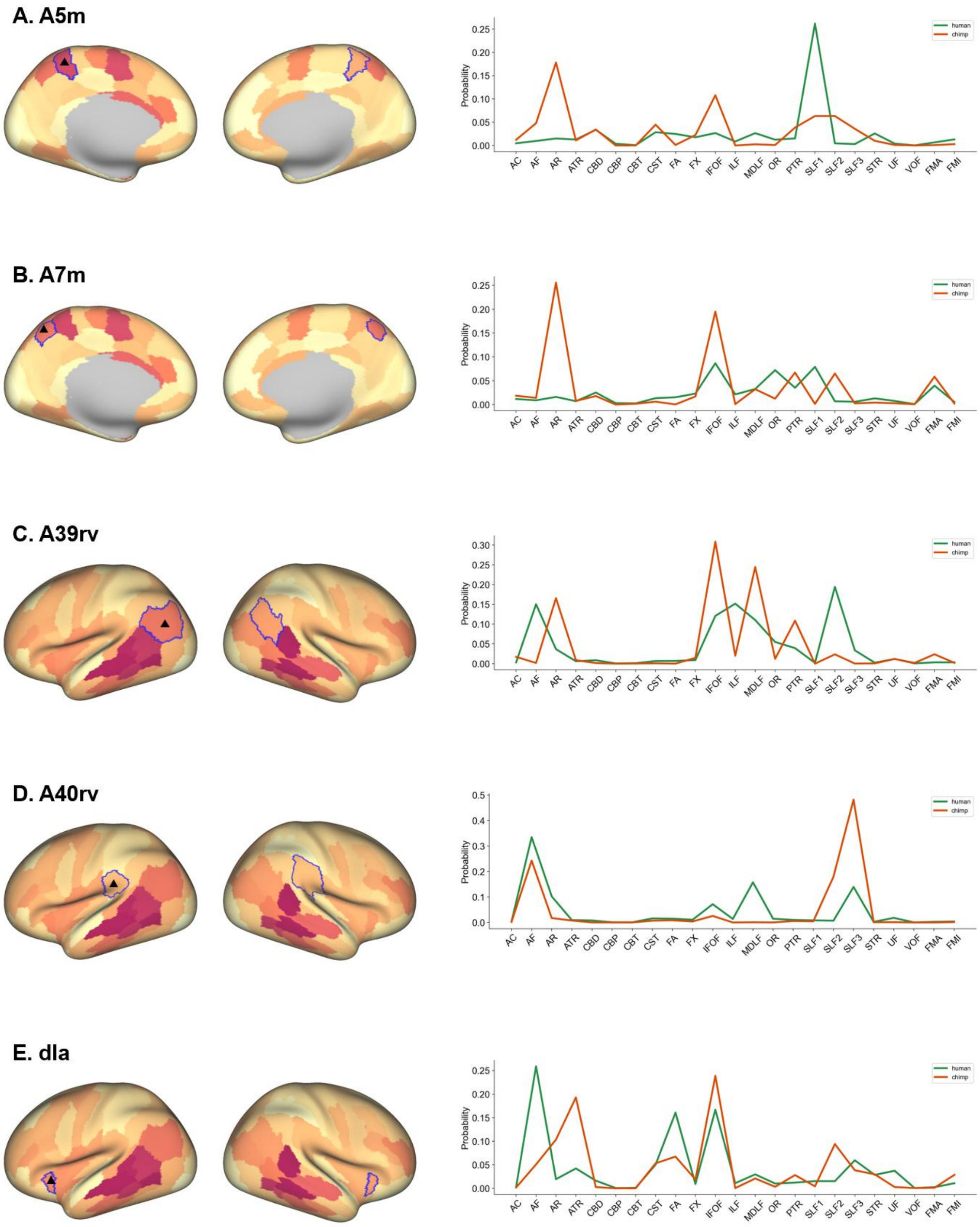


**Figure S38. Tract connectivity distribution of example subregions corresponding to subregions with higher connectivity divergence in Figure 3C. (A)** Tract connectivity distribution of the A5m. **(B)** Tract connectivity distribution of the A7m. **(C)** Tract connectivity distribution of the A39rv. **(D)** Tract connectivity distribution of the A40rv. **(E)** Tract connectivity distribution of the dIa. A5m, medial area 5; A7m, medial area 7; A39rv, rostroventral area 39; A40rv, rostroventral area 40; dIa, dorsal agranular insula.

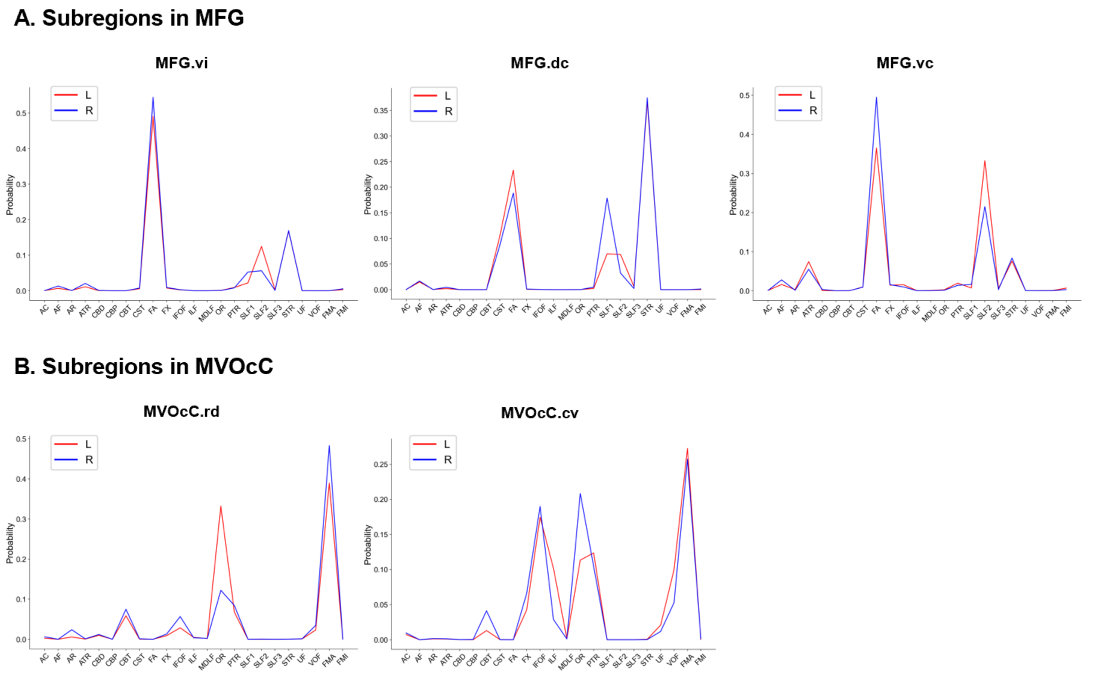


**Figure S39. Tract connectivity distribution of example subregions. (A)** Tract connectivity distribution of subregions in the MFG. **(B)** Tract connectivity distribution of subregions in the MVOcC. MFG.vi, middle frontal gyrus, ventrointermediate part; MFG.dc, middle frontal gyrus, dorsocaudal part; MFG.vc, middle frontal gyrus, ventrocaudal part; MVOcC.rd, medioventral occipital cortex, rostrodorsal part; MVOcC.cv, medioventral occipital cortex, caudoventral part.

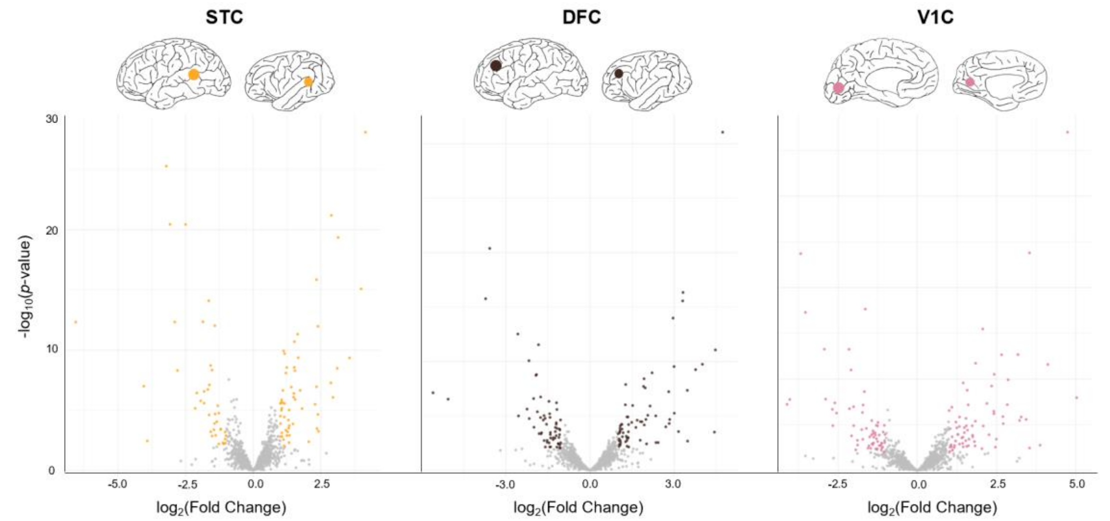


**Figure S40. The differentially expressed genes between humans and chimpanzees for each region.** The differentially expressed genes were calculated using DESeq and were called for FDR < 0.01. 122 genes were found in STC, 92 genes in DFC, and 118 genes in V1C. STC, superior temporal cortex; DFC, dorsolateral frontal cortex; V1C, primary visual cortex.

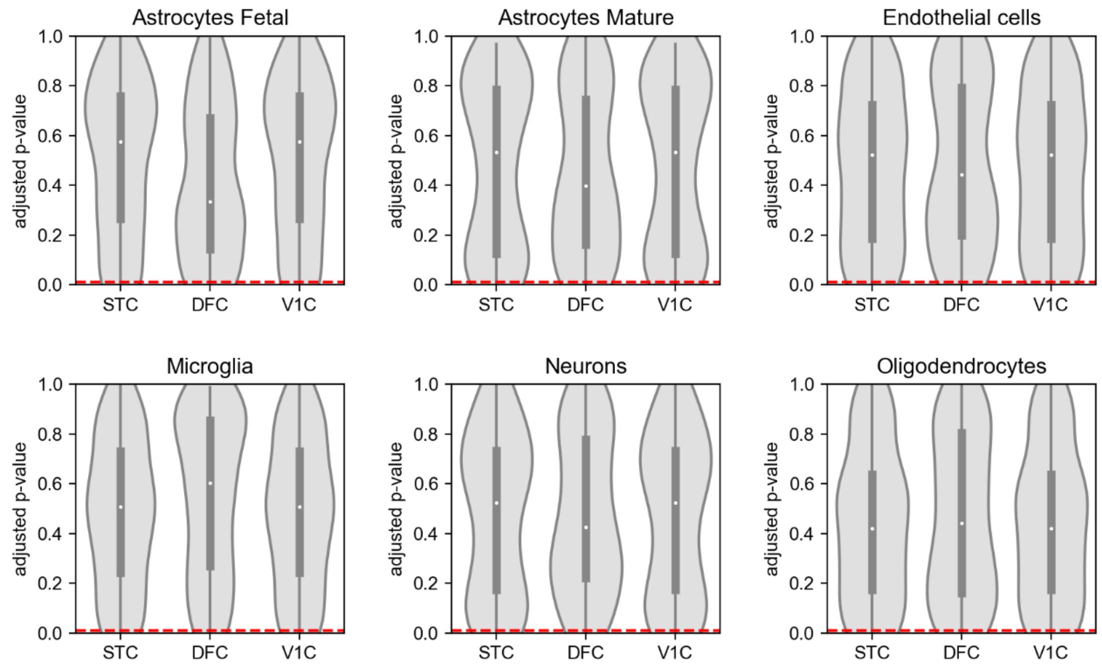


**Figure S41. The differential gene expression analysis was not attributable to variations in the ratio of major cell types between species.** The distribution of adjusted *p*-values (FDR) of cell-type enriched genes in differential gene expression analysis. The dot in the middle denotes the median. The red line shows the cutoff for significance in our differential gene expression analysis (FDR < .01).
